# The spinal cord facilitates cerebellar upper limb motor learning and control; inputs from neuromusculoskeletal simulation

**DOI:** 10.1101/2023.03.08.531839

**Authors:** Alice Bruel, Ignacio Abadía, Thibault Collin, Icare Sakr, Henri Lorach, Niceto R. Luque, Eduardo Ros, Auke Ijspeert

## Abstract

Complex interactions between brain regions and the spinal cord (SC) govern body motion, which is ultimately driven by muscle activation. Motor planning or learning are mainly conducted at higher brain regions, whilst the SC acts as a brain-muscle gateway and as a motor control centre providing fast reflexes and muscle activity regulation. Thus, higher brain areas need to cope with the SC as an inherent and evolutionary older part of the body dynamics. Here, we address the question of how SC dynamics affects motor learning within the cerebellum; in particular, does the SC facilitate cerebellar motor learning or constitute a biological constraint? We provide an exploratory framework by integrating biologically plausible cerebellar and SC computational models in a musculoskeletal upper limb control loop. The cerebellar model, equipped with the main form of cerebellar plasticity, provides motor adaptation; whilst the SC model implements stretch reflex and reciprocal inhibition between antagonist muscles. The resulting spino-cerebellar model is tested performing a set of upper limb motor tasks, including external perturbation studies. A cerebellar model, lacking the implemented SC model and directly controlling the simulated muscles, was also tested in the same benchmark. The performances of the spino-cerebellar and cerebellar models were then compared, thus allowing directly addressing the SC influence on cerebellar motor adaptation and learning, and on handling external motor perturbations. Performance was assessed in both joint and muscle space, and compared with kinematic and EMG recordings from healthy participants. The differences in cerebellar synaptic adaptation between both models were also studied. We conclude that the SC facilitates cerebellar motor learning; when the SC circuits are in the loop, faster convergence in motor learning is achieved with simpler cerebellar synaptic weight distributions. The SC is also found to improve robustness against external perturbations, by better reproducing and modulating muscle cocontraction patterns.

**Summary:** Accurate motor control emerges from complex interactions between different brain areas, the spinal cord (SC), and the musculoskeletal system. These different actors contribute with distributed, integrative and complementary roles yet to be fully elucidated. To further study and hypothesise about such interactions, neuromechanical modelling and computational simulation constitute powerful tools. Here, we focus on the SC influence on motor learning in the cerebellum, an issue that has drawn little attention so far; does the SC facilitate or hinder cerebellar motor learning? To address this question, we integrate biologically plausible computational models of the cerebellum and SC, equipped with motor learning capability and fast reflex responses respectively. The resulting spino-cerebellar model is used to control a simulated musculoskeletal upper limb performing a set of motor tasks involving two degrees of freedom. Moreover, we use kinematic and EMG recordings from healthy participants to validate the model performance. The SC fast control primitives operating in muscle space are shown to facilitate cerebellar motor learning, both in terms of kinematics and synaptic adaptation. This, to the best of our knowledge, is the first time to be shown. The SC also modulates muscle cocontraction, improving the robustness against external motor perturbations.

## 1 Introduction

Accurate motor control enables interactions with the environment and others, a process in which sensory information is integrated by the central nervous system (CNS) and translated into muscle activity, eventually driving body motion. Body motion results from the interaction between the musculoskeletal system and diverse neural regions with distributed, integrative and complementary roles [1]. In the brain, various neural regions project descending motor control signals to the spinal cord (SC); e.g., the motor cortex, involved in the volitional control of motion [2]; the basal ganglia, involved in selecting motor behaviour and balance control [3, 4]; the cerebellum, involved in motor coordination and learning [5]. The SC circuits integrate those motor descending signals to regulate motoneuron activity, ultimately driving muscle activation. Besides, the SC also implements its own motor control mechanisms; e.g., fast reflexes, control of rhythmic locomotion movements, or responses against perturbations [5].

Motor control within the CNS could be synthesised as a hierarchical process; higher brain areas govern motor functions such as planning or learning, and the SC then integrates their descending control signals, provides faster and lower-level control mechanisms, and ultimately drives muscle activity. To comprehend and hypothesise about this hierarchical interaction, neuromechanical modelling and computational simulation represent powerful tools, providing a holistic view conjugating from neuron to neural network to motor behaviour levels [6]. To that aim, we present a hierarchical structure comprising: a cerebellar model, a higher brain area equipped with motor learning and adaptation; an SC model, integrating the cerebellar descending control signals and implementing fast-reflexes and muscle activity regulation, and finally actuating a musculoskeletal upper limb model. This spino-cerebellar integration thus provides a computational exploratory framework, which was further complemented with kinematic and EMG data validation. Both the cerebellum and SC main physiological mechanisms have been previously described, however, little attention has been put on the SC influence on cerebellar motor control. Spinal circuits are evolutionary old, they were present in the first vertebrates emerged about 500 million years ago [7] and fully allowed basic locomotion [8]. As new higher neural areas evolved to handle more complex motor control, they had to coexist and interact with the old lower spinal circuits. It is not clear whether that interaction facilitates motor control or implies a constraint with which higher neural regions have to live with. On the one hand, the SC benefits motor control providing fast feedback loops, lower dimensionality for planning and control, and motor primitives (i.e., low level motor building blocks). On the other hand, higher brain areas have to deal with the highly non-uniform control space and hidden states in the SC, and the need for inverse models that cover not only the body dynamics but also the SC dynamics. Here, we study whether the SC facilitates cerebellar motor learning, or it is simply an evolutionary constraint to be handled.

The cerebellum is key in motor control and coordination, and most importantly motor learning [9]. The Marr-Albus-Ito theory on cerebellar function [10] established the computational principles for supervised cerebellar learning [11], by which the cerebellum enables the adaptation of our actions so their consequences match up to our expectations, i.e., minimising the difference between our intention and the actual movement [12]. This motor learning capability stands upon the plasticity exhibited at the synapses from parallel fibres (PF), i.e., axons of granule cells (GC), to Purkinje cells (PC); plasticity regulated by the action of climbing fibres (CF) reaching PCs [13]. The Marr-Albus-Ito theory assumes the GCs carry a recoding of the sensory inputs conveyed through mossy fibres (MF) [14], whereas CFs carry an instructive signal coding the disparity between our motor expectation and the actual motor state. Despite the well-accepted common ground on the cerebellum established by the Marr-Albus-Ito theory, new findings keep refining the understanding about cerebellar structure and operation, for which computational models are key contributors [15]. Computational models of the cerebellum have been used to study its inner dynamics [16, 17], as well as harnessing cerebellar motor adaptation capabilities to develop adaptive controllers based on internal model building [18, 19].

Lower down in the CNS hierarchy, the SC transmits control signals from brain motor areas to the muscles, and it also conveys sensory signals from muscle receptors back to the brain. But its role in motor control goes beyond a mere gateway between the brain and muscles [20, 21]. The SC contains neural pathways that regulate muscle activity, control reflex responses and produce rhythmic locomotion movements. These spinal pathways channel the sensory feedback mainly from stretch sensitive muscle spindles and tension sensitive Golgi tendon organs (GTO). This sensory feedback is then transmitted to motoneurons through afferent fibres and spinal interneurons, allowing reflex responses and muscle regulation mechanisms: e.g., stretch velocity reflex, static stretch reflex, Golgi tendon reflex, or reciprocal inhibition between antagonist muscles [5]. Besides, these spinal pathways are modulated by higher brain areas during movement execution such as between the stance and swing phases during gait [22, 23], or during arm movements [24, 25, 26], thus highlighting the importance of the interaction between the SC and higher brain areas.

Computational models have been used to gain deeper insight on the SC role in motor control; e.g., control of centre-out reaching movements [27]; control of biceps stretch reflex [28]; reflex modulation via feedback gains [29]; rejection of dynamic perturbations, highlighting the latency hierarchy levels of feedback [30], or the contribution of GTO feedbacks [31]. However, these approaches lacked complex descending signals from higher brain areas, usually applying open-loop supraspinal modules, hence hindering their use to study the interaction between the SC and higher neural regions; larger scale models are required.

Little work has been done on large scale modelling to dig into the SC interaction with higher CNS regions. A recent example coupled spinal circuits with sensory and motor cortex models, forming a feedback control loop designed to reduce the difference between the desired and perceived state of a planar six-muscle arm [32]. The model showed motor control success and reproduced some previous experimental phenomena, whilst it was suggested that the ataxic nature of the produced movements could be due to the lack of a cerebellum model in the loop.

Regarding spino-cerebellar integration in particular, a few previous computational approaches exist. Contreras-Vidal et al. modelled a cerebellum cooperating with an SC-based muscular force model, together with a central pattern generator representing the motor cortex and basal ganglia [33]. The cerebellar model, developed in analogue form and lacking the temporal correlation nature of cerebellar learning, succeeded in learning muscle synergies, including cocontraction of antagonist pairs, that improved upon the SC feedback control of tracking. Different cerebellar lesions were studied, but the influence of the SC in cerebellar motor adaptation was sidestepped. Subsequently, Spoelstra et al. integrated a cerebellar model with an SC model for postural control of a six-muscle two-dimensional arm model [34]. The study assessed the predictive role of the cerebellum in accurate motor control, but again, the effect of the SC in cerebellar learning was not addressed. More recently, Jo integrated a functional cerebellar model with spinal circuits equipped with plasticity but lacking reflex or other complex spinal dynamics [35]. Results showed the effectiveness of the model to learn movements, with synaptic plasticity at the SC helping to acquire muscle synergies. However, as stated by the author, that learning capacity provided to the SC could be located anywhere in the corticospinal pathway, hence loosening possible conclusions on the cerebellum-SC relation.

With the present work, we intend to extend the spino-cerebellar integration studies; we addressed the questions of whether the SC facilitates cerebellar learning or it is just as an evolutionary constraint, and how the SC contributes to handling motor perturbations. We modelled a biologically plausible cerebellar spiking neural network (SNN), equipped with synaptic plasticity at GC-PC connections guided by the instructive signal conveyed through CFs, thus, able to provide motor adaptation. We added an SC model equipped with stretch reflex and reciprocal inhibition, integrating the descending signals from the cerebellum and sending muscle excitation commands to the musculoskeletal upper limb model, equipped with two degrees of freedom (DOF) actuated by eight Hill-based muscles. Both the cerebellar and SC model were integrated in a negative feedback control loop. The study, developed using computational tools and neuromechanical modelling, is also supported by lab recorded kinematics and EMG data from healthy participants.

In the presented framework, the cerebellar model provides the motor adaptation required for the musculoskeletal upper limb model to achieve a set of goal motor behaviours, i.e., different upper limb movements are defined in joint space (position and velocity), and the cerebellum acquires the inverse model allowing accurate position and velocity tracking. We suggest the SC fast control primitives and regulation of muscle activity to be key in facilitating the cerebellar learning of the muscle dynamics; the SC allowed faster motor learning with simpler cerebellar synaptic adaptation. We also hypothesise that the SC plays a major motor control role through cocontraction modulation; i.e., regulation of simultaneous activation of antagonist muscles. Cocontraction has been shown to improve stability by increasing joint apparent stiffness [36], enhance upper limb movement accuracy [37], and it has also appeared useful in movements requiring robustness against perturbations [38]. We found that the stretch reflex and reciprocal inhibition mechanisms participate in modulating cocontraction, with a significant impact on cerebellar motor adaptation and response against external perturbations.

## 2 Results

We integrated the spinal cord and cerebellum models in an upper limb musculoskeletal feedback control loop (Fig. 1A). The spino-cerebellar model commanded the upper limb to perform a set of motor tasks, a motor benchmark divided in two groups: i) lab recorded upper limb movements performed by two healthy participants to study natural self-selected movements, ii) lab designed upper limb movements with bell-shaped velocity profiles to study standard characteristic reaching movements. A cerebellar model lacking the SC integration performed in the same motor benchmark (Fig. 1B) thus providing a spino-cerebellar vs. cerebellar control framework that allowed contextualising the SC and cerebellum integration (see Methods for a further description of the control loop and motor benchmark).

**Fig. 1.**
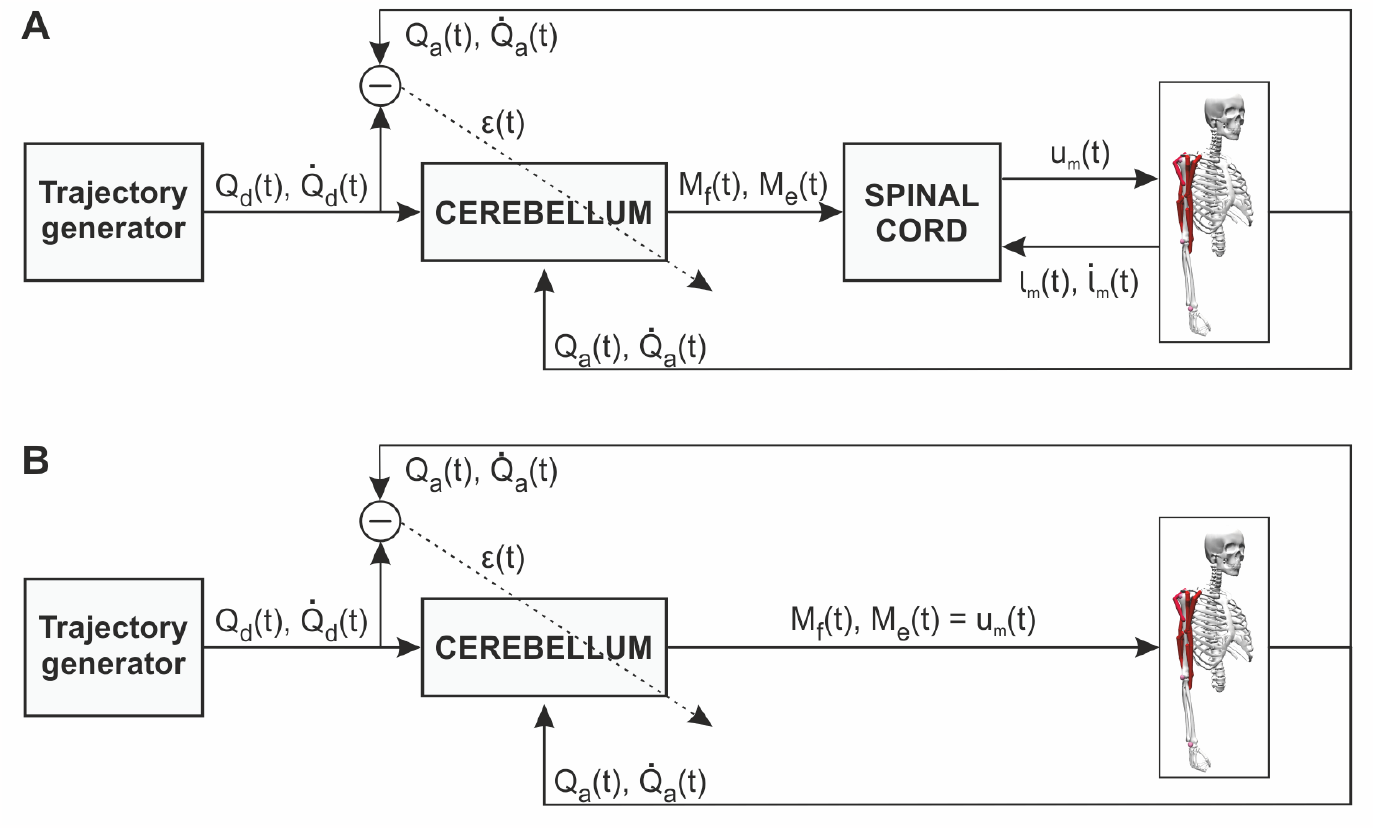
Spino-cerebellar and cerebellar control loops. **A)** Spino-cerebellar model. The cerebellum received the following input sensory information: the desired trajectory (joint position, *Q*_*d*_, and velocity, 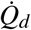) coming from a trajectory generator, representing the motor cortex and other motor areas performing motor planning and inverse kinematics; the actual upper limb state (joint position, *Q*_*a*_, and velocity,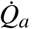) received from the musculoskeletal model; the instructive signal (*ϵ*) obtained as the mismatch between the desired and actual joint state. The cerebellum then generated two output control signals per joint (*M*_*f*_ and *M*_*e*_, for flexor and extensor muscles, respectively), which were processed at the spinal cord. The spinal cord also received the muscle state (length, *l*_*m*_, and velocity,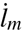) and generated the final muscle excitation signals (*u*_*m*_) which actuated the musculoskeletal model. The musculoskeletal model included two joints, shoulder and elbow, actuated by eight muscles: deltoid posterior and biceps long as shoulder flexors; deltoid anterior and triceps long as shoulder extensors; biceps long, short and brachialis as elbow flexors; triceps long, lateral and medial as elbow extensors. **B)** Cerebellar model. *M*_*f*_ and *M*_*e*_ were directly applied as muscle excitation signals commanded to the upper limb. For bi-articular muscles (biceps long and triceps long), the resulting *u*_*m*_ was the mean of the control signal (*M*_*f*_ or *M*_*e*_) from both joints.

The following sections present the validation of the spino-cerebellar model with the lab recorded kinematics and EMG data; an evaluation of the SC effect in cerebellar motor adaptation in joint, synaptic and muscle spaces; and testing the response against external motor perturbations.

### 2.1 Spino-cerebellar and cerebellar models perform the recorded kinematics

We extracted kinematics and EMG data from two healthy participants (P1 and P2) performing upper limb movements in the vertical plane involving the shoulder and elbow (see Methods). The motor tasks performed by P1 and P2 can be grouped in: i) flexion-extension movements, ii) hand-tracked circular trajectories. Both motor task groups were performed at different speeds, thus providing a set of natural upper limb trajectories which constituted our initial motor control benchmark. We used the joint kinematics (i.e., shoulder and elbow position, *Q*_*d*_, and velocity, 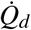) extracted from the recording sessions as the desired trajectory to be learnt by the spino-cerebellar (Fig. 1A) and cerebellar (Fig. 1B) models in the simulation framework. Both models performed 2000 consecutive trials of each desired trajectory, a trial-and-error process that allowed motor adaptation to fully deploy from scratch. The performance metric was given by the position and velocity mean absolute error (MAE), i.e., difference between the desired and actual trajectory in joint space, allowing to assess motor behaviour (see Methods).

We first calculated the position and velocity MAE evolution for both the spino-cerebellar and cerebellar models performing the trajectories extracted from each participant (Fig. 2A, P1’s 1.8s circle trajectory; Fig. 3A, P2’s 1.2s flexion-extension; see Supporting Information for all P1 and P2 motor tasks MAE evolution (S1A to S9A Fig.)). As the trajectory was repeated over time, the cerebellar adaptation allowed position and velocity error reduction. At the end of the motor adaptation process, both the spinocerebellar and cerebellar models followed the target kinematics (Fig. 2B and Fig. 3B; see Supporting Information for all P1 and P2 motor tasks kinematics performance (S1B,C to S9B,C Fig.)).

**Fig. 2.**
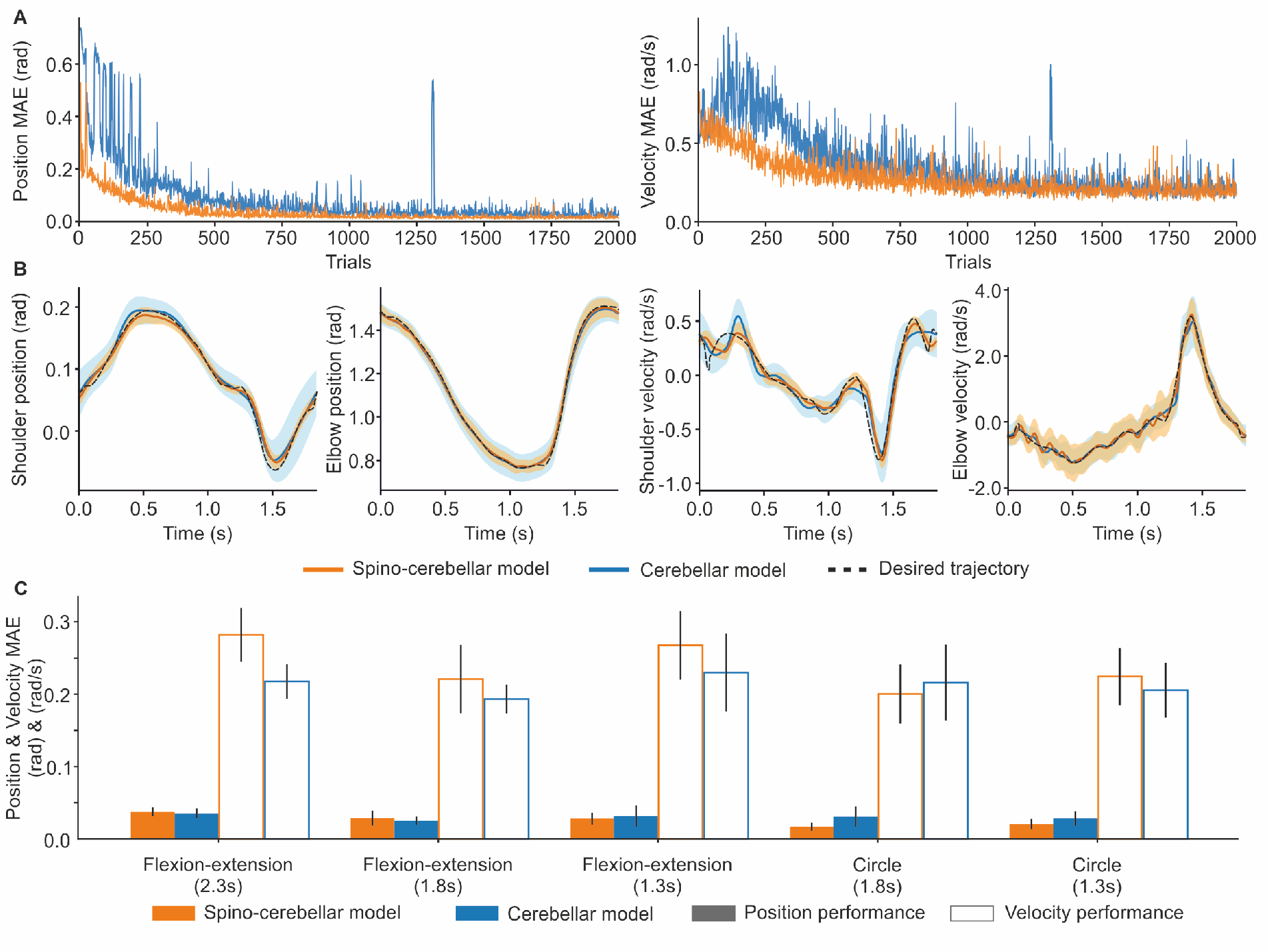
Spino-cerebellar and cerebellar models kinematics performance for the lab recorded scenario, participant 1 (P1). **A)** Position and velocity mean absolute error (MAE) over the 2000-trial motor adaptation process for both the spino-cerebellar and cerebellar models performing P1’s slow circle trajectory (1.8s). **B)** Joint kinematics of the last 200 trials (mean and *std*) for both models performing P1’s slow circle trajectory (1.8s). **C)** Mean and *std* of the position and velocity MAE (last 200 trials) for all P1 recorded trajectories. All spino-cerebellar vs. cerebellar mean position and velocity MAE have a T-test p-value ≤ 0.001.

**Fig. 3.**
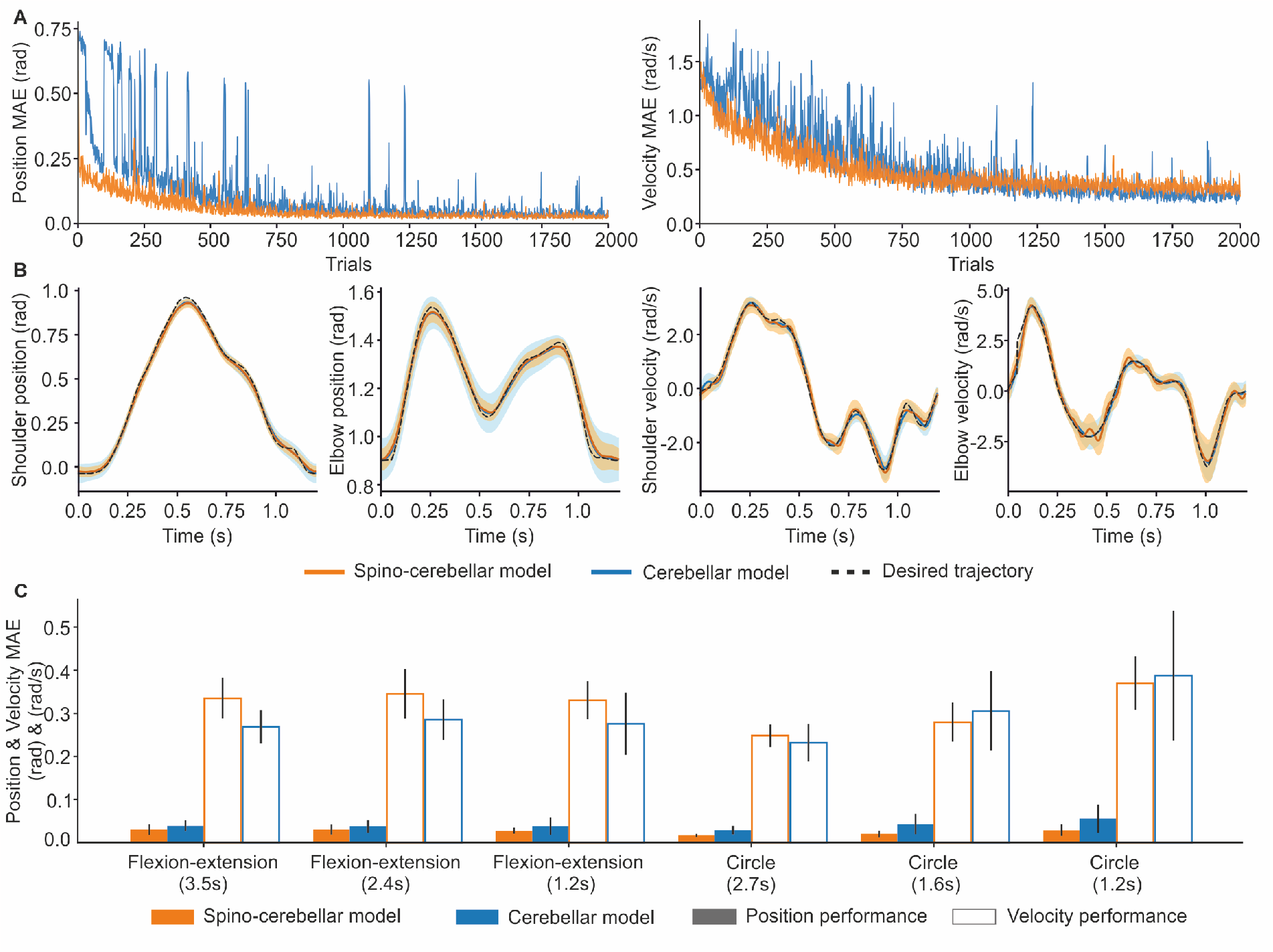
Spino-cerebellar and cerebellar models kinematics performance for the lab recorded scenario, participant 2 (P2). **A)** Position and velocity mean absolute error (MAE) over the 2000-trial motor adaptation process for both the spino-cerebellar and cerebellar models performing P2’s fast flexion-extension (1.2s). **B)** Joint kinematics of the last 200 trials (mean and *std*) for both models performing P2’s fast flexion-extension (1.2s). **C)** Mean and *std* of the position and velocity MAE (last 200 trials) for all P2 recorded trajectories. All spino-cerebellar vs. cerebellar mean position and velocity MAE have a T-test p-value ≤ 0.001, except for fast circle (1.3s) velocity with a p-value = 0.136.

We found that, attending to the MAE mean and standard deviation (std) of the last 200 trials of the motor adaptation process (Fig. 2C and Fig. 3C), the spino-cerebellar model reached better performance in terms of position tracking for all trajectories except for P1’s slow (2.3s) and moderate (1.8s) flexionextension (all spino-cerebellar vs. cerebellar mean position MAE having a T-test p-value ≤ 0.001). Conversely, the cerebellar model reached better performance in terms of velocity tracking except for P1’s slow (1.8s) circle and P2’s moderate (1.6s) and fast (1.2s) circle (all spino-cerebellar vs. cerebellar mean velocity MAE having a T-test p-value ≤ 0.001, except for P2’s fast circle (1.2s) with a p-value = 0.136).

### 2.2 The spinal cord improves cerebellar learning convergence and speed

Once we revealed the adaptation capability of both the spino-cerebellar and cerebellar models, we studied the influence of the SC model on cerebellar learning over the adaptation process. Using the position and velocity MAE evolution of each P1 and P2 trajectory, we compared the spino-cerebellar and cerebellar models learning convergence and learning speed. To study learning convergence we applied control charts on the MAE data to determine the number of trials required to achieve a stable performance [39]. To check learning speed we analysed the number of trials required for the mean MAE of 200 samples to reach a given target (i.e., 0.1 rad for position MAE, and 0.5 rad/s for velocity MAE). Learning convergence and speed were tested on both position and velocity tracking performance (see Fig. 4A for an example of position and velocity MAE evolution and the metrics used, see Methods for a further description).

**Fig. 4.**
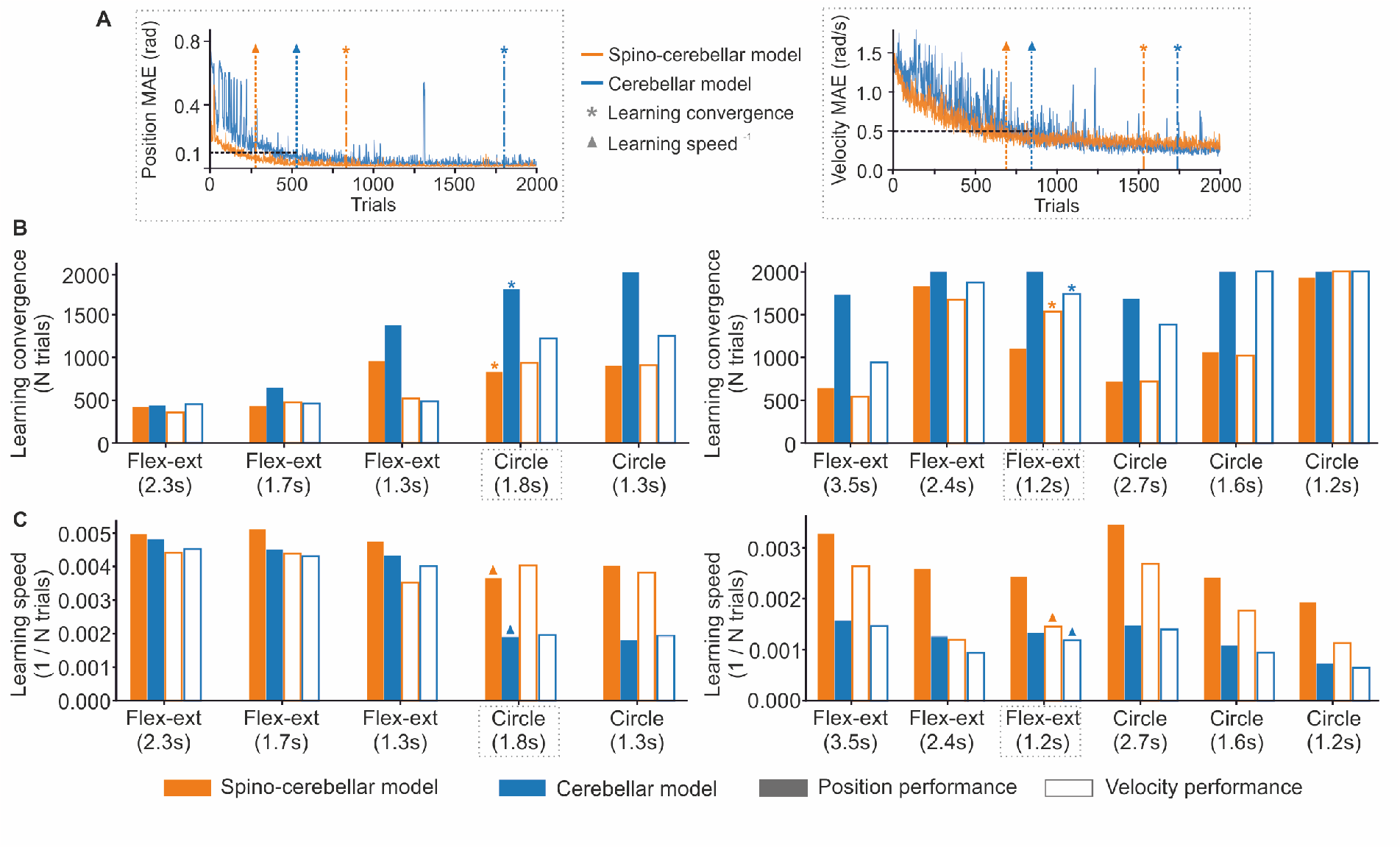
Spino-cerebellar and cerebellar models motor adaptation for all P1 and P2 recorded trajectories. **A)** Position MAE for the spino-cerebellar and cerebellar models for P1 slow circle (left column), and velocity MAE for both models performing P2 fast flexion-extension (right column). Both MAE plots show the trials at which the learning convergence and learning speed metrics are fulfilled. **B)** Learning convergence for both models and all trajectories from P1 (left column) and P2 (right column). The bar plots display the number of trials required by each model to fulfill the learning convergence criteria (see Methods). **C)** Learning speed for both models and all trajectories from P1 (left column) and P2 (right column). The bar plots depict the inverse of the number of trials required to reach a position MAE of 0.1 rad and a velocity MAE of 0.5 rad/s.

The SC was proven to facilitate cerebellar learning as it reduced learning convergence time (Fig. 4B), and increased learning speed (Fig. 4C) for both position and velocity for both P1 and P2 trajectories (Fig. 4 left and right column, respectively). Thus, cerebellar motor adaptation was shown to be: i) stabilised by the SC: average convergence time for *MAE*_*pos*_ was 988 ± 466 trials for the spino-cerebellar model, and 1607 ± 536 trials for the cerebellar model; and for *MAE*_*vel*_ 978 ± 512 trials, and 1255 ± 581 trials, respectively; ii) accelerated by the SC: average learning speed for position was 3.5*e*−3 ± 1.0*e*−3 trials^−1^ for the spino-cerebellar model, and 2.3*e*−3 ± 1.4*e*−3 trials^−1^ for the cerebellar model; and for velocity 2.8*e*−3 ± 1.2*e*−3 trials^−1^, and 2.1*e*−3 ± 1.4*e*−3 trials^−1^, respectively.

### 2.3 The spinal cord simplifies cerebellar synaptic adaptation at GC-PC

Consistently with the Marr-Albus-Ito cerebellar theory, learning in the cerebellum was provided by means of an STDP mechanism adjusting the synaptic weights at GC to PC synapses (a connection established through PFs, i.e., GC axons). The effect of the SC on cerebellar learning, already checked in terms of motor performance in the previous section, must leave its trace at the level of cerebellar synaptic adaptation.

During the motor adaptation process of both the spino-cerebellar and cerebellar models, we recorded the synaptic weight evolution at GC-PC connections every 200 trials for all P1 and P2 trajectories (Fig. 5A and B). We then measured the entropy of the GC-PC synaptic weight distributions to quantify the synaptic complexity for both models: the higher the entropy, the more complex the synaptic weight distribution, i.e., higher heterogeneity of synaptic weights at the GC-PC population. Contrasting the synaptic entropy of both models allowed evaluating the effect of the SC on cerebellar synaptic adaptation (Fig. 5C and D). Noteworthy, results showed that for all motor tasks the SC reduced the entropy of the synaptic weight distribution: the mean entropy over all P1 and P2 trajectories was 3.65 ± 0.78 for the spino-cerebellar model, and 4.41 ± 1.14 for the cerebellar model. When the SC was lacking in the control loop, more complex synaptic patterns (i.e., higher specialisation) were required at cerebellar GC-PC connections. The spino-cerebellar model showed a simpler distribution of synaptic weights at GC-PC connections; in other words, the spinal cord was therefore shown to simplify learning in the cerebellum.

**Fig. 5.**
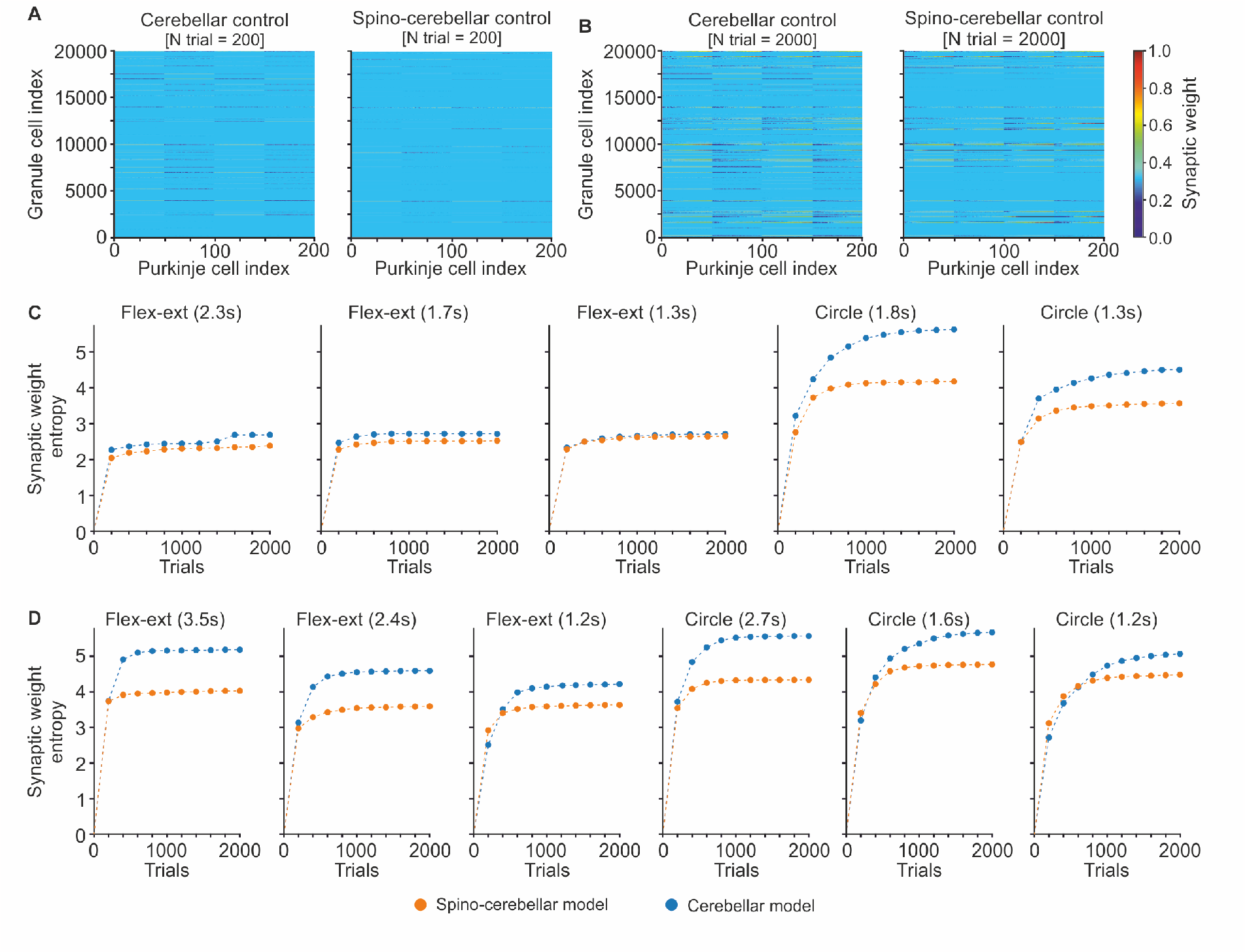
Spino-cerebellar and cerebellar synaptic entropy. **A), B)** Synaptic weights at granule cell - Purkinje cell synapses, after 200 and 2000 trials respectively, for both models performing P1’s 1.8s circle trajectory. The heat map represents the normalised GC-PC synaptic weights, which could range from 0.0 to 15.0 nS. **C), D)** Evolution of the synaptic entropy at the GC-PC synapses over the 2000-trial motor adaptation process, for all P1 and P2 trajectories, respectively. The higher the entropy, the more complex the GC-PC synaptic distribution (i.e., higher heterogeneity in the synaptic weights over the GC-PC synapses).

### 2.4 Spino-cerebellar and cerebellar outcome in muscle space

We then evaluated the outcome in muscle space of both the spino-cerebellar and cerebellar models. We compared the recorded EMG envelopes to the main activated muscles during P1 and P2 trajectories (Fig. 6A, deltoid posterior (DELTpost) and brachialis (BRA) for P1 slow circle, deltoid anterior (DELTant) and triceps lateral (TRIlat) for P2 fast flexion-extension; please find a figure displaying all the recorded EMG in Supporting Information (S13 and S14 Fig.)). Both models reproduced the main activation patterns of each muscle with a small shift for P2 DELTant and TRIlat. The correlation between the spino-cerebellar or cerebellar activation and the EMG signals was generally larger than 0.5 (see Supporting Information (S15 and S16 Fig.)). The correlation was, however, larger for the spino-cerebellar model for most of the muscles and scenarios (all spino-cerebellar vs. cerebellar correlation having a T-test p-value ≤ 0.001). Nevertheless, the correlation averaged over muscles was similar between the two models for all the movements and we could not conclude on a better muscle pattern reproduction by one or the other model (all spino-cerebellar vs. cerebellar mean correlation having a T-test p-value ≥ 0.05).

**Fig. 6.**
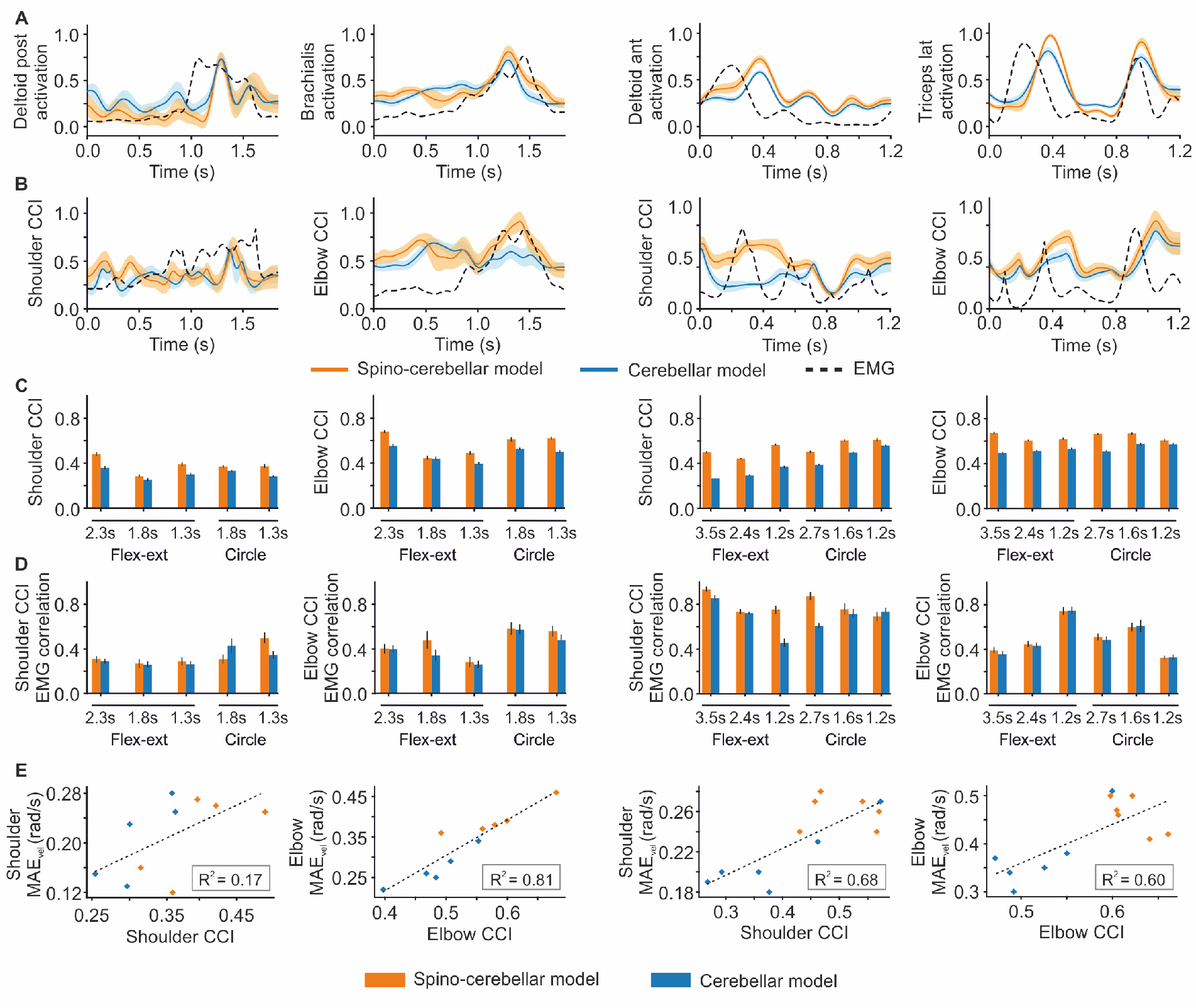
Spino-cerebellar and cerebellar model performance in muscle space for all P1 and P2 recorded trajectories. **A)** Comparison of muscle activation signals with recorded EMGs: the comparison only shows the main activated muscles during recordings of P1’s slow circle (two left columns) and P2’s fast flexion-extension (two right columns). The plots show the muscle activity of the 200 trials prior to reaching the learning convergence metric, as well as their mean and std, for the two models performance. EMG signals are scaled by the maximum of the activation signals for each muscle for the sake of representation. **B)** Joint cocontraction indexes (CCI) from EMG activity and both models performance, for the trajectories represented in A). EMG CCI are scaled by the maximum of the models CCI for the sake of representation. **C)** Joint CCI values for both models and all P1 (two left columns) and P2 (two right columns) trajectories. All spino-cerebellar vs. cerebellar CCI have a T-test p-value ≤ 0.001. **D)** Joint CCI correlation between the models and EMG for all P1 (two left columns) and P2 (two right columns) trajectories. All spino-cerebellar vs. cerebellar CCI correlation have a T-test p-value ≤ 0.001, except for the fast trajectories for the elbow. **E)** CCI-*MAE*_*vel*_ relation: linear regression between joint CCI and joint *MAE*_*vel*_ over all the trajectories from P1 (two left columns) and P2 (two right columns).

Results might not be conclusive when referred to a direct, muscle by muscle comparison between our models performance and the recorded EMG; note that our musculoskeletal upper limb model was actuated by 8 muscles, a mere simplification of the complex muscle dynamics of the human upper limb. To overcome this, we further studied performance in muscle space using the joint cocontraction index (CCI), thus unifying muscle activity per joint and providing a more comprehensive analysis (see Methods). We found that the spino-cerebellar model better reproduced the CCI patterns at the level of the elbow for P1 slow circle and at the level of the shoulder for P2 fast flexion-extension (Fig. 6B). Significantly, the spino-cerebellar model provided a higher CCI for all P1 and P2 trajectories, both for the shoulder and elbow (Fig. 6C, all spino-cerebellar vs. cerebellar CCI having a T-test p-value ≤ 0.001). We then compared the CCI provided by both models with the CCI from the recorded EMG (Fig. 6D). The correlation was mainly higher for the spino-cerebellar model for all the trajectories except P1’s slow circle (1.8s) and P2’s fast circle (1.2s) for the shoulder (all spino-cerebellar vs. cerebellar correlation with a T-test p-value ≤ 0.001, except for the fast trajectories for the elbow). We observed a similar trend as that observed for *MAE*_*vel*_, therefore, we performed a linear regression between the CCI and *MAE*_*vel*_ for each joint. The results (Fig. 6E) highlighted a linear trend between these quantities for P1 elbow and P2 shoulder and elbow (with a coefficient of determination of 0.81, 0.68 and 0.60 respectively), whereas P1 shoulder presented a weaker relationship (with a coefficient of determination of 0.17).

Overall, we highlighted various findings that were consistent for various trajectories with various initial and final positions and speeds. The spino-cerebellar model provided more stable and faster learning with simpler cerebellar synaptic adaptation, and an increase in CCI with better correlation to the recorded EMG.

### 2.5 The spinal cord increases the robustness against motor perturbations

To study the response against external perturbations of both the spino-cerebellar and cerebellar models, we used our lab designed benchmark: upper limb flexion-extension movements with bell-shaped velocity profiles, characteristic of reaching movements [40]. This kind of movement is usually used for addressing active-limb control malfunctioning, as cerebellar patients usually display upper limb oscillatory tremors that result in endpoint overshooting and undershooting when reaching a target [41].

Both models faced 2000 consecutive trials of the flexion-extension movement performed at different speeds (3s, 2.3s, 1.5s); after motor adaptation, both models succeeded in performing the target kinematics (see Supporting Information (S17 Fig.)). Once both models adapted to perform the desired trajectories, we tested the contribution of the SC in handling motor perturbations. For that, we induced a set of external forces: i.e., 50 N for 30 ms applied to the hand in different directions and at different points along the flexion-extension movement, resulting in kinematics deviation (Fig. 7A). We then measured the MAE deviation from the ideal, no-perturbation scenario (Fig. 7B). Each perturbation type was applied on 10 separate trajectory trials to get an average response (see Methods). The cerebellar learning capability was disabled to avoid adaptation to the perturbations. Fig. 7A displays the kinematics performance of both models under one perturbation type, whilst performing the moderate flexion-extension movement (2.3s). Larger kinematic deviation can be observed for the cerebellar model compared to the spino-cerebellar one, especially at the level of the elbow. Fig. 7B presents spino-cerebellar and cerebellar model response against all the perturbations applied during the moderate flexion-extension trajectory. The spino-cerebellar model shows smaller MAE deviation 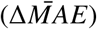 in position performance for all perturbation types, and smaller velocity 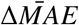 for all perturbations except for the first, fourth and sixth perturbation types. It can be noted that there were both inter and intra-variability in the effect of the various perturbation types on the trajectory kinematics, resulting in T-test p-values ≥ 0.05 for spino-cerebellar vs. cerebellar MAE deviation for some cases (see Fig. 7 caption). Similar results were obtained for the slow and fast bell-shaped flexion-extension trajectories; please find the corresponding figure in Supporting Information (S18 Fig.). The spino-cerebellar model was shown to be more robust against perturbations than the cerebellar model.

**Fig. 7.**
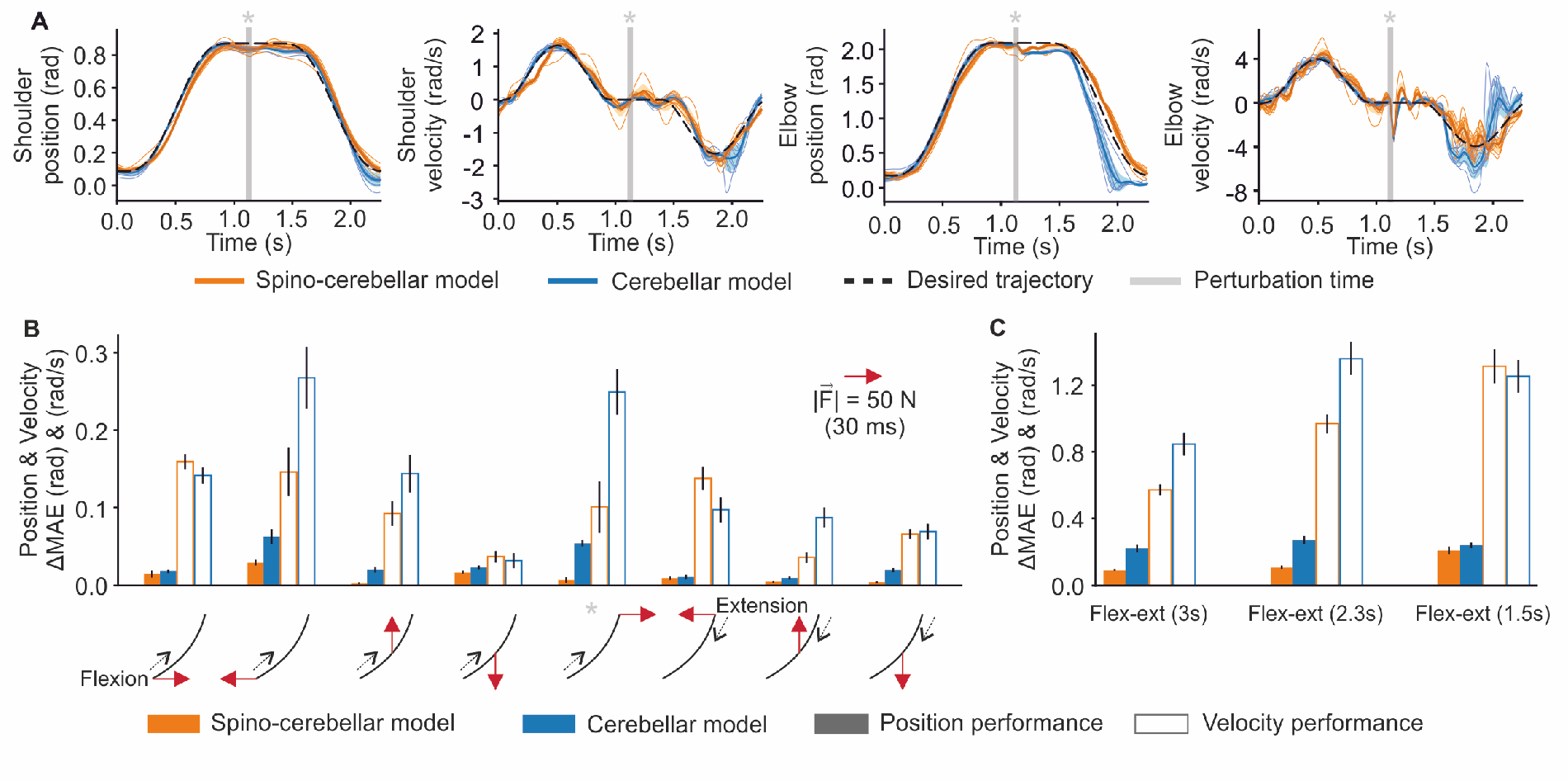
Spino-cerebellar and cerebellar model response against external force perturbations during bell-shaped flexion-extension trajectories. **A)** Kinematics performance for both the spino-cerebellar and cerebellar models under one forward perturbation at the flexed position whilst performing the 2.3s flexion-extension trajectory. 10 trials are displayed. **B)** Position and Velocity MAE deviation 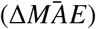 due to all the perturbations applied during the 2.3s flexion-extension trajectory. Mean 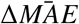 and *se* of 10 trials are displayed. Spino-cerebellar vs. cerebellar MAE deviation have a T-test p-value ≤ 0.05 for position for the 3rd, 5th, 6th, 7th and 8th perturbation type, and for velocity for the 1st, 2nd, 4th, 6th and 7th, whereas the other cases have a T-test p-value ≥ 0.05. **C)** Mean 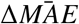 and *se* for all the perturbations applied to the flexion-extension trajectories with different speeds (3s, 2.3s, 1.5s). 10 perturbed trials were used for each perturbation type. All spino-cerebellar vs. cerebellar mean MAE deviations have a T-test p-value ≥ 0.05.

Fig. 7C finally presents the average MAE deviation over all the applied perturbations for the three bell-shaped, flexion-extension trajectories. The spino-cerebellar model results in larger velocity MAE deviation for the fast trajectory, but lower MAE deviation for all the other cases (all spino-cerebellar vs. cerebellar mean MAE deviation having a T-test p-value ≥ 0.05 due to the variable effect of the various perturbation types). The SC is thus shown to help handling motor perturbations in most cases.

## 3 Discussion

The integration of biologically plausible computational models of neural regions allows studying their interaction and complementarity. We presented a computational exploratory approach integrating a cerebellar and an SC model, performing motor control of an upper limb musculoskeletal model; a simulation framework complemented with kinematic and EMG data validation. We contrasted the spino-cerebellar integrated model with a cerebellar model, both performing in the same motor benchmark, which allowed us to extract some key elements of the kinematic and muscle performance directly attributable to the presence of the SC in the spino-cerebellar control loop. The SC was found to stabilise and accelerate cerebellar motor adaptation and to improve the response against perturbations through stretch reflexes and reciprocal inhibition. Rather than being an evolutionary constraint, the SC offers motor control benefits.

Both the spino-cerebellar and cerebellar models succeeded in learning the musculoskeletal dynamics to achieve the goal motor behaviour. Noteworthy, the presence of the SC provided faster motor adaptation, thus assisting cerebellar learning. In this regard, a significant finding was the fact that the spino-cerebellar model revealed less complexity at the GC-PC synaptic weight distribution; i.e., the SC led to the formation of less specialised GC-PC synapses in the cerebellum. To the best of our knowledge, it is the first time that a computational model highlights and weights the influence of the SC in facilitating cerebellar learning. Direct regulation of muscle activity by the SC has been here found to facilitate the cerebellar acquisition of the upper limb inverse dynamics. Indeed, the body plant dynamics to be learnt by higher brain areas, might be simplified by the SC taking over lower level and faster control primitives, such as the SC potential role in gravity compensation [42, 43]. Thus, the SC performance in muscle space may lighten other operations of the sensorimotor process, occurring at a higher level such as the cerebellum’s contribution in compensating interaction torques in joint space [44], or in shaping spatiotemporal muscle synergies rather than generating specific complex muscle patterns [45].

The SC stabilises the system at muscle level, increasing cocontraction through stretch reflexes and coordinating the antagonist activation patterns through reciprocal inhibition. Thus, the SC participates in modulating cocontraction, which plays an important role in motor control and stability [36, 38], providing better accuracy despite its energy cost [37]. In our framework, the spino-cerebellar model increased the joint CCI in all the studied motor tasks compared to the cerebellar model; i.e., cocontraction was indeed mostly determined by the SC motor action. Importantly, the CCI from the spino-cerebellar model also resulted in a better correlation with the CCI patterns from the recorded EMG signals, thus supporting closer biological plausibility than the cerebellar model. The CCI increment was also revealed when inducing perturbations in the control loop; the spino-cerebellar model provided a better response, reducing the kinematic deviation. Muscle elasticity has been previously pointed as a key passive contributor in handling perturbations [46]. Our framework, using the same muscle mechanical properties in both the spino-cerebellar and cerebellar control loops, allowed directly assigning to the SC a pivotal role in providing robustness against external perturbations, thus supporting previous findings [30, 31].

The cocontraction increase carried by the SC involved a poorer velocity tracking. Indeed, the spinal reflexes between antagonist muscles may induce oscillatory activation patterns and thus alter the velocity performance. We did not observe, however, any trend in CCI values related to movement speeds despite higher cocontraction values have been reported in slower movements [38]. Due to the SC and cerebellar models conception, our implementation lacks differentiation between the roles of the cerebellum and SC depending on movement speed. Note however that it is expected a major role of the cerebellum in fast ballistic movements which cannot rely on feedback availability [47, 34], and which do present lower cocontraction levels [38].

Our model could be further improved by adding other cocontraction mechanisms to the control loop. Clinical studies supported a potential role of the cerebellum and basal ganglia in cocontraction mechanisms. In particular, patients with cerebellar ataxia showed excessive agonist-antagonist coactivation [48] and cerebellar stimulation was shown to reduce coactivation in patients with spasticity [49]. Thus, future development of the cerebellar model shall include control of the cocontraction level. On the SC side other pathways could be included, in particular modulation mechanisms that are present during arm movements [24, 25, 26]. For instance, presynaptic inhibition of Ia terminals at both activated and antagonist pathways is slightly decreased at the onset of a voluntary contraction through descending signals. Thus, the increased gain of the stretch reflex pathway ensures that activated motoneurons receive Ia feedback support. The reciprocal Ia inhibition is also depressed during a voluntary contraction at the corresponding muscle to prevent its inhibition by the stretch-induced Ia discharge from its antagonist. During cocontraction, reciprocal Ia inhibition is also depressed by increased presynaptic inhibition on Ia terminals [5]. Also synaptic plasticity could be included in the SC model, as done in previous computational approaches [32]. Activity dependent plasticity mechanisms have been reported in the SC: e.g., the spinal stretch reflex can indeed be conditioned [50]; the feedforward circuits within the SC, in addition to somatosensory feedbacks, may contribute to SC learning by allowing motoneurons to contrast feedforward and feedback motor inputs [51]. Supporting the latter, [52] showed that signals in human muscle spindle afferents during unconstrained wrist and finger movements predict future kinematic states of their parent muscle. Muscle spindles would then have a forward-sensory-model role, as that attributed to the cerebellum [53], emphasising the complementarity and overlapping functionality between neural regions.

Integrated computational models represent a powerful tool to support and guide experimental studies in the pursuit of a better understanding of the CNS. We believe our spino-cerebellar model to contribute in this direction, providing a picture of how the SC influences cerebellar motor adaptation and learning. Further development of the model, together with addition of other neural regions, will help to keep elucidating CNS operation.

## 4 Methods

The cerebellar and SC models operated in a closed loop with joint and muscle feedback (Fig. 1A), in which cerebellar motor learning was assisted by fast reflex response and muscle activity regulation provided by the SC. The cerebellar model received sensory input signals describing the desired motor state (desired position, *Q*_*d*_, and velocity, 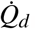, per joint) and the actual motor state of the upper limb musculoskeletal model (actual position, *Q*_*a*_, and velocity, 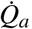, per joint). The comparison of the desired and actual motor states provided the instructive signal (*E* per joint), also received by the cerebellum. The cerebellar output comprised a flexor-extensor (i.e., agonist-antagonist) pair of control signals (*M*_*f*_ and *M*_*e*_ per joint) that were sent to the SC model, which also received direct muscle feedback (length, *l*_*m*_, and velocity, 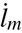, per muscle). The SC generated the muscle excitation signals (*u*_*m*_ per muscle) resulting in muscle activation which finally actuated the upper limb musculoskeletal model, thus closing the loop. To contextualise the spino-cerebellar integration, we also implemented the control loop lacking the SC circuits (Fig. 1B). In this scenario, the cerebellar output signals were directly used as muscle excitation signals. The control loop included sensory and motor delays, mimicking the biological pathways. In the cerebellar sensorimotor pathway, there exists a delay ranging from about 100 to 150ms (with inter and intraindividual variations), accounting for the time spent from the generation of a motor command until sensing back its effect [54]. Regarding the SC to muscles transmission, a delay of about 30ms has been reported for the upper limb [55, 56]. Our spino-cerebellar model included a 50ms sensorial delay affecting the reception of sensory inputs in the cerebellum; a transmission delay of 30ms from the cerebellum to the SC, and 30ms from the SC to the muscles, total motor delay of 60ms. The asymmetry between sensory and motor delay stands for the higher latency found in neuromuscular junction, electromechanical and force generation delays (involved in the motor pathway), compared to the sensing, nerve conduction and synaptic delays (involved in the sensory pathway) [57].

The following subsections describe the different components of our spino-cerebellar control loop. The various building blocks were integrated using Robot Operating System (ROS), allowing a modular implementation.

### 4.1 Cerebellar model

We implemented a spiking neural network (SNN) replicating some cerebellar neural layers and equipped with spike-timing-dependent plasticity (STDP) to allow motor learning and adaptation. The cerebellar SNN model was adapted from previous models applied to robot control loops [18, 19]. The cerebellar SNN structure was divided in the following neural layers: i) mossy fibres (MFs), constituted the sensory input layer conveying the desired and actual motor state signals 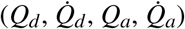; ii) the spiking activity of MFs was transferred through excitatory afferents to the granule cell (GC) layer, where the sensory input information was univocally recoded; iii) the axons of the GCs, i.e., the parallel fibres (PFs), formed excitatory connections with the Purkinje cells (PCs); iv) PCs also received the excitatory action of the climbing fibres (CFs) conveying the instructive signal (*E*); v) the deep cerebellar nuclei (DCN) layer received the inhibitory action from PCs and excitatory connections from both MFs and CFs. The DCN spiking activity was translated into output motor commands (flexor-extensor motor control signals, *M*_*f*_ and *M*_*e*_) that constituted the cerebellar motor response to the sensory stimuli. Every neural layer was divided in two microcomplexes [58], being each microcomplex oriented to drive one of the two joints (shoulder or elbow). Each microcomplex at the PC-CF-DCN loop was partitioned into two regions: agonist and antagonist. The agonist region operated the joint flexor muscles, whereas the antagonist region operated the extensor muscles. This synergic agonist-antagonist (flexor-extensor) architecture allowed the cerebellar model to regulate the spatiotemporal muscle activity patterns [45], key for successful motor control [59]. See Fig. 8 for a schematic representation of the cerebellar network, and Table 1 for network topology.

**Table 1.** Cerebellar neural topology. Dashed entries stand for not applicable.

| Neurons |  | Synapses |  |  |  |
| --- | --- | --- | --- | --- | --- |
| Pre-synaptic | Post-synaptic | Number | Type | Initial Weight (nS) | Weight range (nS) |
| 80 MFs | 20x10<sup>3</sup> GCs | 80x10<sup>3</sup> | AMPA | 0.18 | - |
| 80 MFs | 200 DCN | 16x10<sup>3</sup> | AMPA | 0.3 | - |
| 20x10<sup>3</sup> GCs | 200 PCs | 4000x10<sup>3</sup> | AMPA | 4.8 | [0.0, 15.0] |
| 200 PCs | 200 DCN | 200 | GABA | 1.0 | - |
| 200 CFs | 200 PCs | 200 | AMPA | 0.0 | - |
| 200 CFs | 200 DCN | 200 | AMPA | 0.5 | - |
| 200 CFs | 200 DCN | 200 | NMDA | 0.25 | - |

**Fig. 8.**
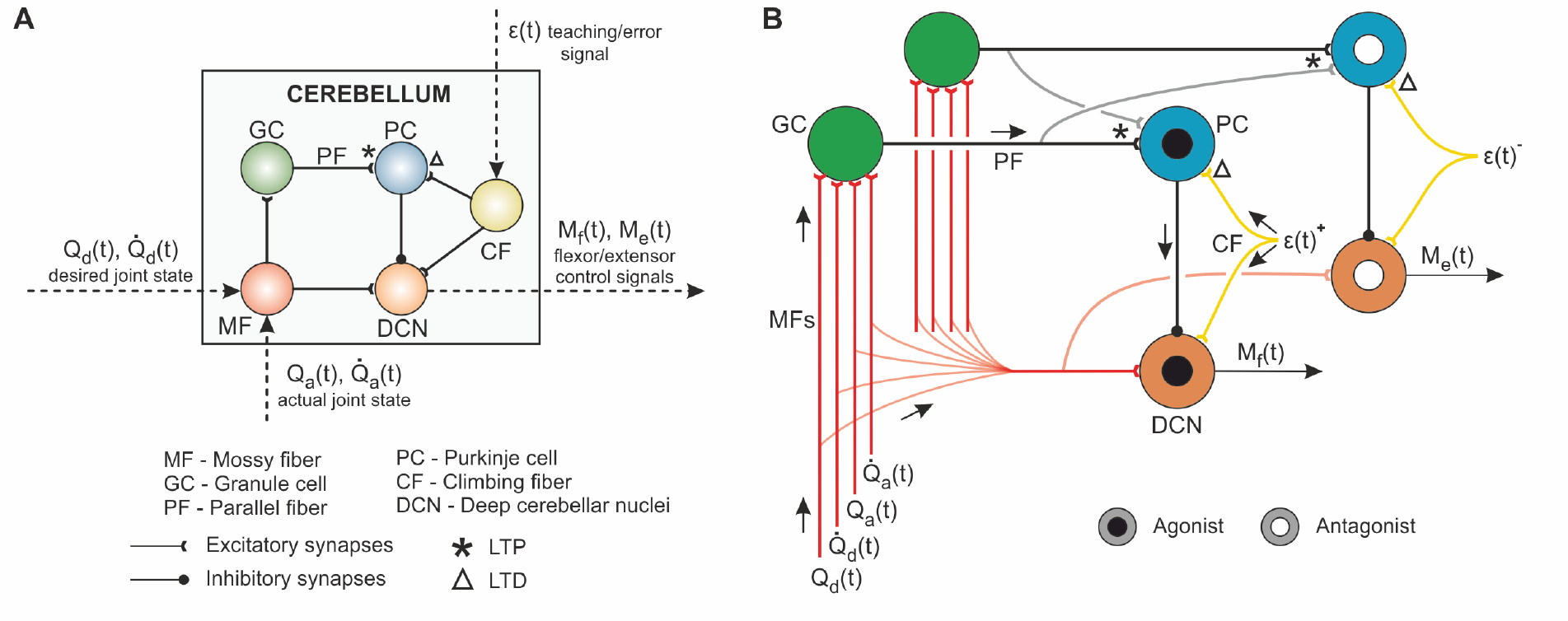
Cerebellum model. **A)** Neural layers, connections, input and output sensorimotor signals. The input signals are conveyed by the mossy fibres (MFs), which project excitatory synapses to the granule cells (GCs). These perform a recoding of the input signals, and project excitatory connections through the parallel fibres (PFs) reaching Purkinje cells (PCs). PF-PC connections are endowed with plasticity, balanced between the long-term potentiation (LTP) caused by the input PF spikes, and longterm depression derived from the climbing fibres (CFs) activity reaching PCs. CFs convey the instructive signal. Finally, PCs project inhibitory synapses towards the deep cerebellar nuclei (DCN), the output layer of the cerebellar model, which also receives a baseline excitatory action from MFs and CFs. **B)** Detailed schematic of the cerebellar connections. Each GC receives the input excitatory action from a unique combination of four MFs. Each input signal 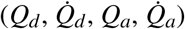, is codified by ten MFs, being only one out of the ten MFs active at each time step. Hence, at each time step, four MFs will be active (one per input signal). That unique combination of four input MFs excites one single GC, allowing to perform a univocal representation of the sensory input at the granular layer. PCs then receive the excitatory action from all GCs in the cerebellar model and only one CF, allowing to relate the joint-specific instructive signal, to the global sensory state received from GCs. The PC-CF-DCN loop differentiates between agonist and antagonist regions, thus allowing simultaneous control of both flexor and extensor muscles.

Consistently with the Marr-Albus-Ito theory on cerebellar motor adaptation [60, 61, 62], our cerebellar SNN model was equipped with synaptic plasticity at the GC-PC synapses. The synaptic weights were adjusted by means of an STDP mechanism that correlated the sensory information (univocally coded at GCs and transferred to PCs through PFs) and the instructive signal (conveyed to PCs by CFs). This STDP mechanism was a balanced process of long-term potentiation (LTP) and long-term depression (LTD). Each time a PC neuron received a GC spike through a PF, that synapse was potentiated (LTP) by a fixed amount as follows:

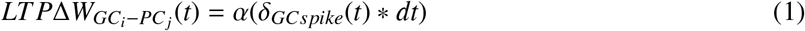

where 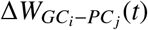 stands for the synaptic weight change between GC *i* and PC *j*; *α* = 0.006*nS* is the synaptic weight increment; and *δ*_*GCspike*_(*t*) is the Dirac delta function of a GC spike, received at PCs through PFs.

When the spiking activity of a CF conveyed an instructive signal to a PC neuron, the GC-PC connection that was involved in that error generation was depressed (LTD) as described by:

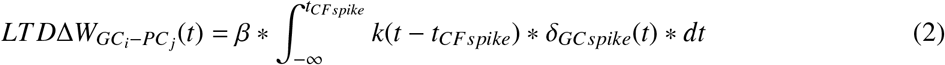

where *β* = −0.003*nS* stands for the synaptic weight decrement; and *k*(*x*) defines the integrative kernel with eligibility trace correlating past sensory inputs with the present instructive signal, i.e., the amount of LTD due to a CF spike depended on the previous GC activity received at PCs through PFs (see [18, 19] for a further description). A well-balanced LTP-LTD process changed the PF-PC synaptic weights, thus modifying the PCs output activity and the inhibitory action of PCs over DCN neurons, which ultimately varied the DCN output activity. Modulating the DCN activity allowed adaptation of the output motor response to the input stimuli. An iterative exposure to the sensory patterns defining the desired motor task, allowed adapting the motor response for error reduction.

We used leaky integrate and fire (LIF) neurons (see Supporting Information (S1)) and EDLUT simulator [63] to build the cerebellar SNN model. Please see [18, 19] for a further review of the STDP mechanism and cerebellar layers.

### 4.2 Spinal cord model

Our SC model integrated the descending control signals from the cerebellum and the direct muscle feedback (Fig. 9A). The SC model allowed fast reflex response and muscle activity regulation by means of monosynaptic Ia stretch reflex and disynaptic reciprocal inhibition pathways between antagonist muscles. The motoneuron (MN) of a given muscle received the following inputs: i) an excitatory connection conveying the cerebellar output signal (*M*_*f*_ or *M*_*e*_, for flexor or extensor muscle); ii) an excitatory connection from the Ia afferent fibre of the muscle (i.e., stretch reflex); iii) an inhibitory connection from the Ia interneuron (Ia IN) innervated by the Ia afferent of the antagonist muscle (i.e., reciprocal inhibition). The antagonist relation between the muscles of the upper limb model is detailed below. The neuron leaky integrate dynamics of the MN firing rate, *r*, were modelled as follows:

**Fig. 9.**
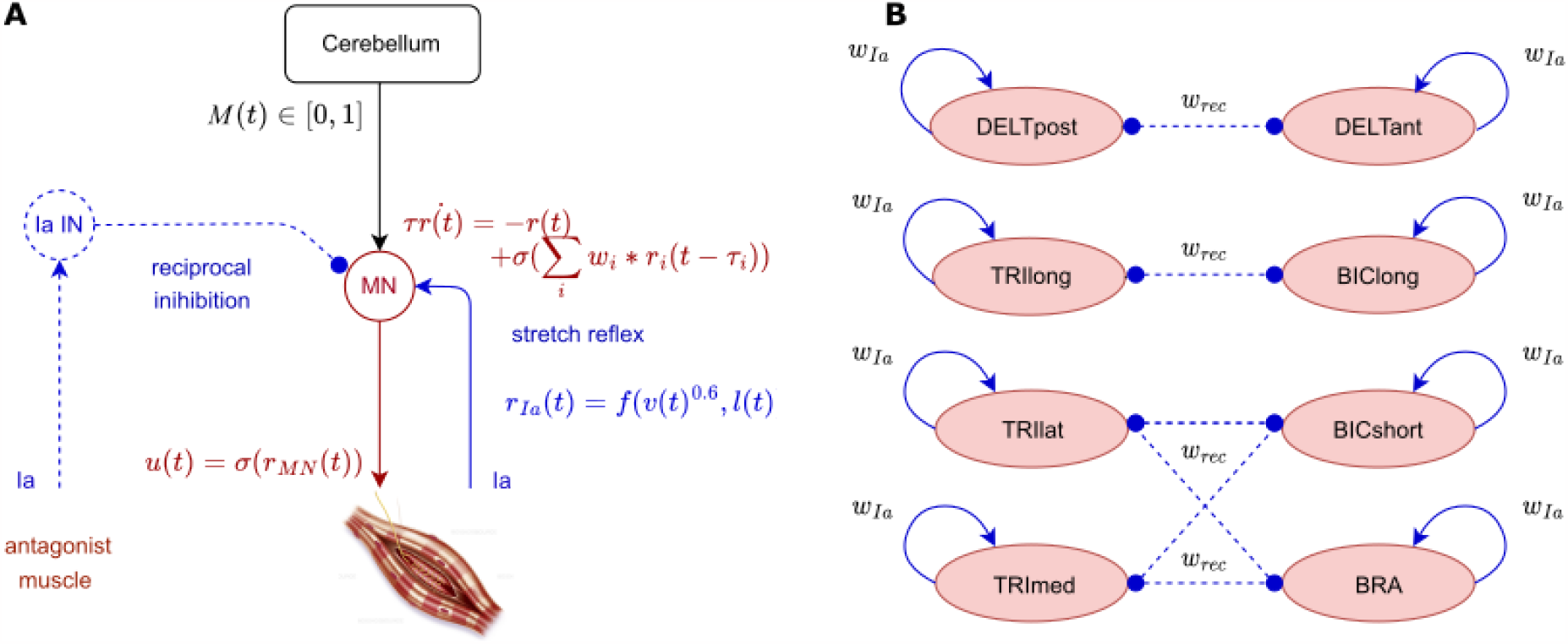
Spinal cord model. **A)** The spinal cord circuits were modelled as one motoneuron per muscle, receiving an excitatory input control signal (M) from the cerebellum, an excitatory connection from the Ia afferent fibre of the muscle (i.e., stretch reflex) and an inhibitory connection from the Ia interneuron (Ia IN) innervated by the Ia afferent of the antagonist muscle (i.e., reciprocal inhibition). We also included inhibitory connections between antagonist Ia interneurons. Each neuron is modelled with leaky integrate dynamics. **B)** Antagonist relation between the 8 upper limb muscles: all the muscles shared the same synaptic weight for the stretch reflex and reciprocal inhibition pathways, i.e., 1 for excitatory synapses and 0.5 for the inhibitory.

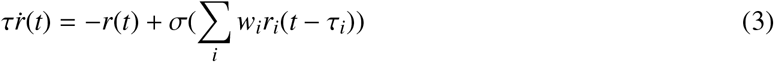

where *τ* = 1ms stands for the spinal neuron activation time constant; 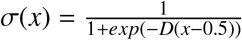 with *D* = 8, emulating the on-off behaviour of neurons; *i* describes the MN input signals; *w*_*i*_ is the synaptic weight of the input connection, being 1 for excitatory synapses and 0.5 for the inhibitory; *r*_*i*_ is the input activity; and *τ*_*i*_ = 30ms stands for the stretch reflex response delay. Depending on neuron size, *τ* can vary from 1 to 10ms [64], we only considered fast-response neurons as in [28]. For the upper limb, *τ*_*i*_ is about 30ms [55, 56]. The output rates of the MNs are finally provided as muscle excitation signals to the musculoskeletal model through a sigmoid (*u*(*t*) = *σ*(*r*(*t*))), thus inducing movement. The dynamics of Ia IN neurons followed the same description, with differing input activity including inhibitory connections between antagonist Ia IN (Fig. 9B).

We used Prochazka’s model for the Ia afferent feedback dynamics [65], with a mean firing rate of 10Hz [28, 66, 67]:

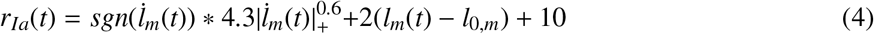

where *l*_*m*_ and 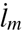 describes the muscle fibre length and velocity in mm and mm/s; and |*x*|_+_= *max*(|*x*|, 0.01). The output rate, *r*_*Ia*_, was scaled by its maximum *r*_*Ia,max*_ to get a normalised value, i.e., *r*_*Ia*_*E*[0, 1].

To model the SC we used FARMS Python library, developed at the BioRobotics laboratory.

### 4.3 Musculoskeletal upper limb model

We used a 2 DOF musculoskeletal upper limb model as the front-end body to be controlled. The model, adapted from [68], included two flexion-extension joints: shoulder and elbow. The model was actuated by 8 Hill-based muscles [69], with the following joint distribution: i) for the shoulder, flexion was carried by the deltoid anterior and posterior (DELTant, DELTpost) and the biceps long (BIClong), and extension was conducted by the triceps long (TRIlong); ii) for the elbow, flexion was provided by the biceps long and short (BICshort) and the brachialis (BRA), whilst extension was allowed by the triceps long, lateral and medial (TRIlat, TRImed). Note that BIClong and TRIlong were bi-articular muscles, as they actuated both joints. The antagonist relation between muscles is depicted in Fig. 9B. The Hill-based muscle dynamics were the following:

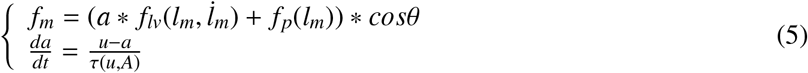

with *f*_*m*_ the muscle force, *f*_*lv*_ a combination of the force-length and force-velocity curves, *f*_*p*_ the passive force-length curve, *θ* the pennation angle, *a* the muscle activation (i.e., the concentration of calcium ions within the muscle), and *u* the muscle excitation (i.e., the firing of the MN) [69]. We used OpenSim physics engine to simulate the muscle and skeleton dynamics [70]. To allow using kinematics and EMG from lab recordings, an OpenSim upper limb model was scaled to match the morphology of each lab participant. This scaling process was achieved using OpenPifPaf Human Pose Estimation algorithm [71] during the static period and OpenSim scaling tool.

### 4.4 Benchmarking with various motor tasks

We used a set of different motor tasks to be performed by the spino-cerebellar and cerebellar models, differentiating between two scenarios: lab recorded and lab designed motor tasks.

For the lab recorded scenario, we used kinematics and EMG recordings from healthy participants performing different arm movements. Experiments were approved by the CER-VD under the license number 2017-02112 and performed in accordance with the Declaration of Helsinki in NeuroRestore laboratory at Lausanne CHUV. Two participants, P1 and P2, were asked to perform planar reaching movements (flexion-extension) and continuous circular movements, both movements performed in the vertical plane and at various speeds (self-selected speeds). For flexion-extension movements both shoulder and elbow moved in the same direction, whilst during the continuous circular movements the joints moved in opposite directions. Thus, our benchmark includes interaction torques both assisting and resisting the movement. The recorded kinematics (i.e., joint position and velocity) constituted the desired motor state (*Q*_*d*_, 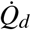) used as the control loop sensory input, whilst the EMG recordings supported model validation in muscle space. For each recorded motor task we ran the experimental setup with both the spino-cerebellar and cerebellar models, using an OpenSim upper limb model scaled to match the participant’s morphology. We then compared the models’ experimental performance to the lab recordings in both joint and muscle spaces.

P1 and P2 movements were recorded using an RGB-D camera, and we used OpenPifPaf human pose estimation algorithm [71] to extract the 2D positions of the participant’s anatomical joints at a frame rate of 25fps. Then 3D pose was deduced from the 2D pose, camera intrinsic, and depth information after accounting for distortion. The occlusions were removed using specially designed filters that ensure coherence in joint anatomy and time. We scaled an OpenSim upper limb musculoskeletal model to match the participant’s morphology, and ran inverse kinematics (IK) over the body segment kinematics, thus allowing the extraction of joint position and velocity from the participant’s motion. P1 generally performed fast movements, and the kinematics recordings of his fast circular movements were too noisy to extract joint position, thus we excluded this scenario from our analysis. For muscle activity, we recorded EMG using Delsys system and Trigno Avanti and Trigno Quattro sensors with a acquisition frequency of 1259.3Hz. We aligned the EMG with the kinematics signals thanks to a trigger inducing a pulse in an additional EMG channel and lightning a led in the camera range. We then computed the EMG envelopes to compare with our models muscle activation signals. For each recorded signal, we removed the mean and rectified the signal, which was then filtered using a low pass Butterworth filter with a 5Hz cutoff frequency. We applied the same processing steps to the maximal voluntary contraction (MVC) signal of each muscle (recorded at the beginning of the session), and used the maximal value of the processed MVC to normalise the corresponding muscle processed EMG signal.

For the lab designed scenario, we implemented a set of flexion-extension movements with different bell-shaped joint velocity profiles, characteristic of multi-joint arm reaching movements [40]. We then used the joint kinematics (*Q*_*d*_, 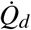) as the desired motor state to be performed by the spino-cerebellar and cerebellar models (please see Supporting Information for a depiction of the bell-shaped trajectories (S10 to S12 Fig.)). We broadened the benchmark by adding a perturbation study using these bell-shaped trajectories. After cerebellar learning consolidation, we applied a set of motor perturbations whilst the trajectories were being performed: 50N for 30ms, applied to the hand in different directions and at different points along the flexion-extension movement. Each perturbation type was applied to 10 separate trials to get an average response, leaving 3 non-perturbed trials in between perturbed trials so that the model returned to its unperturbed state. Note that cerebellar learning was disabled during the perturbation study, to avoid cerebellar adaptation to the external forces and focus on SC response.

Using this motor benchmark, and comparing the performance of the spino-cerebellar and cerebellar models, we could evaluate the cerebellum and spinal cord integration in terms of: muscle activity, motor adaptation and joint space performance, synaptic adaptation, and response to motor perturbations, for various trajectories with different initial and final positions and speeds. Please see Supporting Information for a representation of the motor tasks joint kinematics (S1 to S12 Fig.).

### 4.5 Cerebellar instructive signal

The cerebellar instructive signal *ϵ*(*t*) was obtained as the mismatch between the desired and actual joint state, combining in a single value per joint both position and velocity errors as follows:

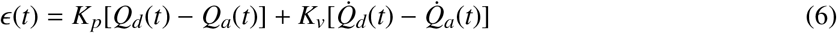

where *K*_*p*_ = 3 and *K*_*v*_ = 1 are the position and velocity error gain, respectively. The trajectory error signal in joint space can be derived from the proprioceptive and sensory information conveyed by the spino-cerebellar tract from the muscle spindles (muscle length) and Golgi tendon organs (muscle force) to the cerebellum [72].

### 4.6 Performance metrics

#### 4.6.1 Measuring kinematics performance

To evaluate the kinematic performance of the spino-cerebellar and cerebellar models, we defined a set of metrics based on the mean absolute error (MAE) between the desired (*Q*_*d*_, 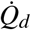) and actual (*Q*_*a*_, 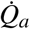) motor state of the arm:

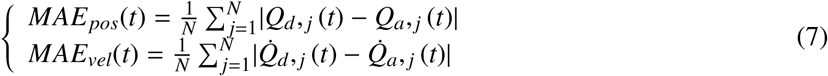

where *N* stands for the number of joints (2), and *j* for the joint index. We considered the position and velocity MAE of each motor task trial to assess the performance accuracy:

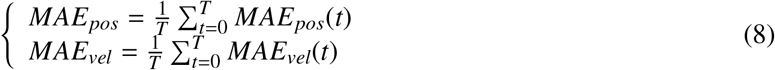

where *T* stands for the motor task period. We finally averaged these values over 200 trials and compared the final performance of the two models with the final mean *MAE*_*pos*_ and 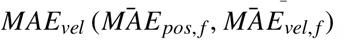. We also computed the standard deviation (*std*) and the T-test p-value between the two models’ results with a T-test for the means of two independent samples of values.

#### 4.6.2 Measuring learning performance

To measure the learning convergence (i.e., number of trials required to reach a stable trajectory tracking), we used control chart metrics [39]. Throughout the *MAE*_*pos*_ and *MAE*_*vel*_ curve of each motor task (all performed for a total of 2000 learning trials) we computed the mean (*μ*) and standard deviation (*σ*) using a sample size of 200 trials, which provided the following performance stability limits:

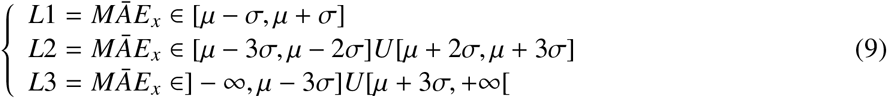

We then checked the percentage of those 200 trials within each limit. As the limits were defined by the *std*, we also checked that the *std* value was below 0.012rad for position and 0.055rad/s for velocity. Thus, at trial *x*, the behaviour was stable if the percentage of the 200 previous trials within each limit fulfilled the metrics defined in Table 2, and the *std* was equal or below the aforementioned values. By comparing the learning convergence of the spino-cerebellar and cerebellar models (i.e., number of trials required to reach a stable performance) we quantified the effect of the SC in the cerebellar motor adaptation process.

**Table 2.** *MAE* convergence criteria from control chart

| Stability limit | $MAE_{pos}$ | $MAE_{vel}$ |
| --- | --- | --- |
| $L1 = \bar{MAE}_x \in [\mu - \sigma, \mu + \sigma]$ | $\geq 78\%$ | $\geq 73\%$ |
| $L2 = \bar{MAE}_x \in [\mu - 3\sigma, \mu - 2\sigma] \cup [\mu + 2\sigma, \mu + 3\sigma]$ | $\leq 3\%$ | $\leq 3\%$ |
| $L3 = \bar{MAE}_x \in ]-\infty, \mu - 3\sigma] \cup [\mu + 3\sigma, +\infty[$ | $\leq 2\%$ | $\leq 2\%$ |
| $\sigma$ | $\leq 0.012$ | $\leq 0.055$ |

Additionally, we assessed the learning speed of the two models by considering the number of trials required to reach a target *MAE*_*pos*_ of 0.1rad and a target *MAE*_*vel*_ of 0.5rad/s. We defined the learning speed metric as 1 over this number of trials 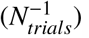.

Thus, we evaluated how long it took for the performance to stabilise (learning convergence) and how fast the performance approached accurate tracking (learning velocity).

#### 4.6.3 Measuring cerebellar synaptic adaptation

To study the effect of the SC in cerebellar synaptic adaptation we quantified the difference in the synaptic weight distribution at GC-PC connections between the spino-cerebellar and cerebellar models. Each PC was innervated by all GCs in the model; i.e., a GC formed an excitatory synapse with each PC (total number of GCs in the model *i* = 20000; total number of PCs in the model *j* = 200). We stored the synaptic weight of all GC-PC synapses in a matrix of size *i*x *j*:

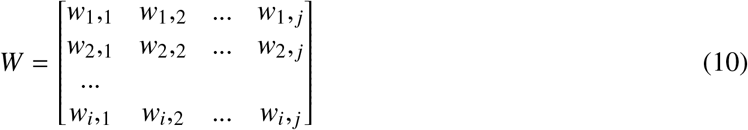

where *w*_*x*_,_*y*_ is the synaptic weight of the synapse between GC *x* and PC *y*.

We then represented the normalised weights stored in W, using *i* as the y-axis and *j* as the x-axis, providing a visual representation of the synaptic weight distribution (Fig. 5). To analyse the differences between the synaptic patterns that were formed in each model, we applied to the images Shannon’s entropy from [73], thus providing a quantitative measure of the complexity of the synaptic distribution. The higher the entropy, the more heterogeneous the synaptic weights; i.e., more specialised GC-PC connections were formed.

#### 4.6.4 Measuring robustness against perturbations

To assess the robustness against perturbations, for each applied perturbation type we computed the mean MAE deviation from the no-perturbation scenario over the 10 perturbed trials as follow:

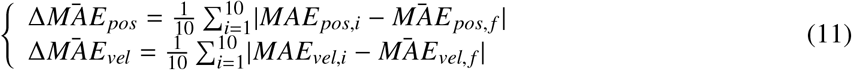

where *MAE*_*x,i*_ is the MAE resulting from the *i*^*th*^ perturbed trial and 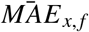 the final MAE for the corresponding no-perturbation scenario. We also computed the *std* and T-test p-value between the spinocerebellar and cerebellar model results as above.

#### 4.6.5 Measuring muscle space performance

We also evaluated performance in the muscle space using the lab recorded benchmark. Activation signals from models are commonly compared to EMG envelopes, but such comparisons are generally difficult to achieve due to scaling issues that hinder a direct analogy between the model and the real muscle dynamics. To overcome this issue, we followed a more comprehensive approach by computing the correlation between activation signals and EMG envelopes. We computed the EMG envelopes by rectifying and low pass filtering the signals using a 5th order Butterworth filter with a cut-off frequency of 5Hz. We also recorded the maximal velocity contraction (MVC) signals for each participant, we processed them the same way and finally normalised the EMG signals by the maximum of the muscle MVC signal. Then, for each movement type, we considered only the main activated muscles with clear activation patterns during the recordings, i.e., DELTant, BIClong, BICshort, TRIlat and BRA for P1 flexion-extension movements; DELTant, DELTpost, BIClong, TRIlat and BRA for P1 circular movements; DELTant, BIClong, TRIlong and TRIlat for P2 flexion-extension movements; and DELTant, DELTpost, and BRA for P2 circular movements. Thus, there is inter-participant variability in muscle patterns. A figure per participant displays all the recorded EMG and highlights these main patterns in Supporting Information (S13 and S14 Fig.). It is worth noting that P1 performed smaller shoulder flexion with larger elbow flexion during flexion-extension movements compared to P2, corresponding to additional BICshort and BRA activation without TRIlong activation. In our experimental setup, we computed the maximal correlation around lag 0 (on a window of one-fourth of the movement duration) for the 200 trials prior to reaching the learning convergence metric and extracted the mean, *std* and T-test p-value between the spino-cerebellar and cerebellar model results. Regarding the lab recorded data, we did not consider those muscles that presented low and noisy EMG signals; however, those muscles were actually activated in our experimental simulations. Our musculoskeletal model indeed contained only 8 muscles, so that such overactivation may reproduce other non-modelled muscle recruitment.

To study our cocontraction hypothesis, we computed and compared the cocontraction index (CCI) for each joint. From lab recordings or experimental simulations, we considered the average of EMG envelop or muscle activation signals, respectively, within each agonist and antagonist muscle group (i.e, DELTant and BIClong for shoulder flexor muscles; DELTpost and TRIlong for shoulder extensors; BIClong, BICshort and BRA for elbow flexors; TRIlong and TRIlat for elbow extensors) and the CCI defined by [74] and assessed by [75]:

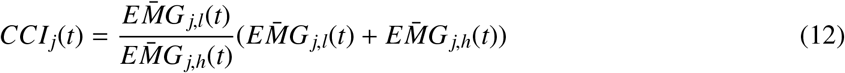

where 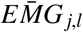 is the level of activity in the less active muscle group and 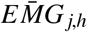 the level of activity in the most active muscle group for each joint. As this index is also sensitive to scaling, we computed the maximal correlation around lag 0 (on a window of one-fourth of the movement duration) for the first 200 trials reaching our learning convergence metric (see Methods) and extracted the mean, *std* and T-test p-value between the spino-cerebellar and cerebellar model results. We also computed the mean joint CCI over each trajectory. A similar trend as that seen for the *MAE*_*vel*_ was observed. We studied this potential relationship through a linear regression over all P1 and P2 trajectories.

## Data availability

For reproducibility and comparative purposes, the source code is available on Zenodo at https://doi.org/10.5281/zenodo.7701436

### Acknowledgments

We gratefully thank the participants for their patience and willingness to collaborate in the recording sessions.

## Additional information

### Funding

This work was supported by European Union Human Brain Project Specific Grant Agreement 3 (H2020-RIA. 945539), by the Spanish Ministry of Science and Innovation to N.R.L. SPIKEAGE [MICINN-PID2020-113422G A-I00] ref [MCIN/AEI/10.13039/501100011033] and to N.R.L. DLROB [TED2021-

131294B-I00] funded by MCIN/AEI/10.13039/501100011033 and by “European Union NextGenerationEU/PRTR”.

### Author contributions

Conceptualization: Alice Bruel, Ignacio Abadía, Eduardo Ros, Niceto R. Luque, Auke Ijspeert

Data Curation: Alice Bruel, Ignacio Abadía

Formal Analysis: Alice Bruel, Ignacio Abadía

Funding Acquisition: Eduardo Ros, Niceto R. Luque, Auke Ijspeert

Investigation: Alice Bruel, Ignacio Abadía, Thibault Collin, Icare Sakr

Methodology: Alice Bruel, Ignacio Abadía

Software: Alice Bruel, Ignacio Abadía

Supervision: Henri Lorach, Eduardo Ros, Niceto R. Luque, Auke Ijspeert

Writing – Original Draft Preparation: Alice Bruel, Ignacio Abadía

Writing – Review & Editing: Alice Bruel, Ignacio Abadía, Thibault Collin, Icare Sakr, Henri Lorach,

Eduardo Ros, Niceto R. Luque, Auke Ijspeert

No competing interests declared by none of the authors.

### Ethics

Experiments were approved by the CER-VD under the license number 2017-02112 and performed in accordance with the Declaration of Helsinki.

## Notes

### Competing Interest Statement

The authors have declared no competing interest.

## References

[1] Rossignol, S., Dubuc, R., and Gossard, J.P. Dynamic sensorimotor interactions in locomotion. Physiological Reviews, 86(1):89–154, January 2006. doi: 10.1152/physrev.00028.2005.

[2] Ebbesen, C.L. and Brecht, M. Motor cortex — to act or not to act? Nature Reviews Neuroscience, 18(11):694–705, October 2017. doi: 10.1038/nrn.2017.119.

[3] Groenewegen, H.J. The basal ganglia and motor control. Neural Plasticity, 10(1-2):107–120, 2003. doi: 10.1155/np.2003.107.

[4] Visser, J.E. and Bloem, B.R. Role of the basal ganglia in balance control. Neural Plasticity, 12 (2-3):161–174, 2005. doi: 10.1155/np.2005.161.

[5] Pierrot-Deseilligny, E. and Burke, D. The circuitry of the human spinal cord: spinal and corticospinal mechanisms of movement. Cambridge University Press, 2012.

[6] Edwards. Neuromechanical simulation. Frontiers in Behavioral Neuroscience, 2010. doi: 10.3389/fnbeh.2010.00040.

[7] Sherman, S.M. and Usrey, W.M. Cortical control of behavior and attention from an evolutionary perspective. Neuron, 109(19):3048–3054, October 2021. doi: 10.1016/j.neuron.2021.06.021.

[8] Grillner, S. Evolution of the vertebrate motor system—from forebrain to spinal cord. Current Opinion in Neurobiology, 71:11–18, 2021. doi: 10.1016/j.conb.2021.07.016.

[9] Ito, M. Mechanisms of motor learning in the cerebellum. Brain research, 886(1-2):237–245, 2000. doi: 10.1016/S0006-8993(00)03142-5.

[10] Kawato, M., Ohmae, S., Hoang, H., and Sanger, T. 50 years since the marr, ito, and albus models of the cerebellum. Neuroscience, 462:151–174, May 2021. doi: 10.1016/j.neuroscience.2020.06.019.

[11] Raymond, J.L. and Medina, J.F. Computational principles of supervised learning in the cerebellum. Annual Review of Neuroscience, 41(1):233–253, July 2018. doi: 10.1146/annurev-neuro-080317-061948.

[12] Medina, J.F. Teaching the cerebellum about reward. Nature Neuroscience, 22(6):846–848, May 2019. doi: 10.1038/s41593-019-0409-0.

[13] Ito, M. and Kano, M. Long-lasting depression of parallel fiber-purkinje cell transmission induced by conjunctive stimulation of parallel fibers and climbing fibers in the cerebellar cortex. Neuroscience letters, 33(3):253–258, 1982. doi: 10.1016/0304-3940(82)90380-9.

[14] Carrillo, R.R., Naveros, F., Ros, E., and Luque, N.R. A metric for evaluating neural input representation in supervised learning networks. Frontiers in Neuroscience, 12:913, 2018. doi: 10.3389/fnins.2018.00913.

[15] Dean, P., Porrill, J., Ekerot, C.F., and Jörntell, H. The cerebellar microcircuit as an adaptive filter: experimental and computational evidence. Nature Reviews Neuroscience, 11(1):30–43, December 2009. doi: 10.1038/nrn2756.

[16] Luque, N.R., Naveros, F., Carrillo, R.R., Ros, E., and Arleo, A. Spike burst-pause dynamics of purkinje cells regulate sensorimotor adaptation. PLOS Computational Biology, 15(3):e1006298, March 2019. doi: 10.1371/journal.pcbi.1006298.

[17] Luque, N.R., Naveros, F., Abadía, I., Ros, E., and Arleo, A. Electrical coupling regulated by GABAergic nucleo-olivary afferent fibres facilitates cerebellar sensory–motor adaptation. Neural Networks, 155:422–438, November 2022. doi: 10.1016/j.neunet.2022.08.020.

[18] Abadía, I., Naveros, F., Garrido, J.A., Ros, E., and Luque, N.R. On robot compliance: A cerebellar control approach. IEEE Transactions on Cybernetics, 51(5):2476–2489, May 2021. doi: 10.1109/tcyb.2019.2945498.

[19] Abadía, I., Naveros, F., Ros, E., Carrillo, R.R., and Luque, N.R. A cerebellar-based solution to the nondeterministic time delay problem in robotic control. Science Robotics, 6(58), September 2021. doi: 10.1126/scirobotics.abf2756.

[20] Brown, T.G. The intrinsic factors in the act of progression in the mammal. Proceedings of the Royal Society of London. Series B, containing papers of a biological character, 84(572):308–319, 1911. doi: 10.1098/rspb.1911.0077.

[21] Weiler, J., Gribble, P.L., and Pruszynski, J.A. Spinal stretch reflexes support efficient hand control. Nature Neuroscience, 22(4):529–533, February 2019. doi: 10.1038/s41593-019-0336-0.

[22] Prochazka, A. Sensorimotor gain control: A basic strategy of motor systems? Progress in Neurobiology, 33(4):281–307, January 1989. doi: 10.1016/0301-0082(89)90004-x.

[23] Büschges, A. and Manira, A.E. Sensory pathways and their modulation in the control of loco-motion. Current Opinion in Neurobiology, 8(6):733–739, December 1998. doi: 10.1016/s0959-4388(98)80115-3.

[24] Fink, A.J.P., Croce, K.R., Huang, Z.J., Abbott, L.F., Jessell, T.M., and Azim, E. Presynaptic inhibition of spinal sensory feedback ensures smooth movement. Nature, 509(7498):43–48, April 2014. doi: 10.1038/nature13276.

[25] Bennett, D.J. Stretch reflex responses in the human elbow joint during a voluntary movement. The Journal of Physiology, 474(2):339–351, January 1994. doi: 10.1113/jphysiol.1994.sp020026.

[26] Shemmell, J., Krutky, M.A., and Perreault, E.J. Stretch sensitive reflexes as an adaptive mechanism for maintaining limb stability. Clinical Neurophysiology, 121(10):1680–1689, October 2010. doi: 10.1016/j.clinph.2010.02.166.

[27] Tsianos, G.A., Goodner, J., and Loeb, G.E. Useful properties of spinal circuits for learning and performing planar reaches. Journal of Neural Engineering, 11(5):056006, August 2014. doi: 10.1088/1741-2560/11/5/056006.

[28] Sreenivasa, M., Ayusawa, K., and Nakamura, Y. Modeling and identification of a realistic spiking neural network and musculoskeletal model of the human arm, and an application to the stretch reflex. IEEE Transactions on Neural Systems and Rehabilitation Engineering, 24(5):591–602, May 2016. doi: 10.1109/tnsre.2015.2478858.

[29] Stienen, A.H.A., Schouten, A.C., Schuurmans, J., and van der Helm, F.C.T. Analysis of reflex modulation with a biologically realistic neural network. Journal of Computational Neuroscience, 23(3), May 2007. doi: 10.1007/s10827-007-0037-7.

[30] Stollenmaier, K., Ilg, W., and Haüfle, D.F.B. Predicting perturbed human arm movements in a neuro-musculoskeletal model to investigate the muscular force response. Frontiers in Bioengineering and Biotechnology, 8, April 2020. doi: 10.3389/fbioe.2020.00308.

[31] Kistemaker, D.A., Soest, A.J.K.V., Wong, J.D., Kurtzer, I., and Gribble, P.L. Control of position and movement is simplified by combined muscle spindle and golgi tendon organ feedback. Journal of Neurophysiology, 109(4):1126–1139, February 2013. doi: 10.1152/jn.00751.2012.

[32] Verduzco-Flores, S.O. and de Schutter, E. Self-configuring feedback loops for sensorimotor control. eLife, 11, November 2022. doi: 10.7554/elife.77216.

[33] Contreras-Vidal, J.L., Grossberg, S., and Bullock, D. A neural model of cerebellar learning for arm movement control: cortico-spino-cerebellar dynamics. Learning and Memory, 3(6):475–502, 1997. doi: 10.1101/lm.3.6.475.

[34] Spoelstra, J., Schweighofer, N., and Arbib, M.A. Cerebellar learning of accurate predictive control for fast-reaching movements. Biological Cybernetics, 82(4):321–333, March 2000. doi: 10.1007/s004220050586.

[35] Jo, S. A computational neuromusculoskeletal model of human arm movements. International Journal of Control, Automation and Systems, 9(5):913–923, October 2011. doi: 10.1007/s12555-011-0512-9.

[36] Latash, M.L. Muscle coactivation: definitions, mechanisms, and functions. Journal of Neurophysiology, 120(1):88–104, July 2018. doi: 10.1152/jn.00084.2018.

[37] Gribble, P.L., Mullin, L.I., Cothros, N., and Mattar, A. Role of cocontraction in arm movement accuracy. Journal of Neurophysiology, 89(5):2396–2405, May 2003. doi: 10.1152/jn.01020.2002.

[38] Disselhorst-Klug, C., Schmitz-Rode, T., and Rau, G. Surface electromyography and muscle force: Limits in sEMG–force relationship and new approaches for applications. Clinical Biomechanics, 24(3):225–235, March 2009. doi: 10.1016/j.clinbiomech.2008.08.003.

[39] Roberts, S.W. A comparison of some control chart procedures. Technometrics, 8(3):411–430, August 1966. doi: 10.1080/00401706.1966.10490374.

[40] Flanagan, J.R. and Ostry, D.J. Trajectories of human multi-joint arm movements: Evidence of joint level planning. pages 594–613, 2006. doi: 10.1007/bfb0042544.

[41] Becker, M.I. and Person, A.L. Cerebellar control of reach kinematics for endpoint precision. Neuron, 103(2):335–348, 2019. doi: 10.1016/j.neuron.2019.05.007.

[42] Klauer, C., Schauer, T., Reichenfelser, W., Karner, J., Zwicker, S., Gandolla, M., Ambrosini, E., Ferrante, S., Hack, M., Jedlitschka, A., et al. Feedback control of arm movements using neuro-muscular electrical stimulation (nmes) combined with a lockable, passive exoskeleton for gravity compensation. Frontiers in Neuroscience, 8:262, 2014. doi: 10.3389/fnins.2014.00262.

[43] Ritzmann, R., Krause, A., Freyler, K., and Gollhofer, A. Gravity and neuronal adaptation. Micro-gravity Science and Technology, 29(1):9–18, 2017. doi: 10.1007/s12217-016-9519-4.

[44] Bastian, A.J., Martin, T., Keating, J., and Thach, W. Cerebellar ataxia: abnormal control of interaction torques across multiple joints. Journal of neurophysiology, 76(1):492–509, 1996. doi: 10.1152/jn.1996.76.1.492.

[45] Berger, D.J., Masciullo, M., Molinari, M., Lacquaniti, F., and d’Avella, A. Does the cerebellum shape the spatiotemporal organization of muscle patterns? insights from subjects with cerebellar ataxias. Journal of Neurophysiology, 123(5):1691–1710, May 2020. doi: 10.1152/jn.00657.2018.

[46] Burdet, E., Tee, K.P., Mareels, I., Milner, T.E., Chew, C.M., Franklin, D.W., Osu, R., and Kawato, M. Stability and motor adaptation in human arm movements. Biological Cybernetics, 94(1):20–32, November 2005. doi: 10.1007/s00422-005-0025-9.

[47] Koster, B. Essential tremor and cerebellar dysfunction: abnormal ballistic movements. Journal of Neurology, Neurosurgery and Psychiatry, 73(4):400–405, October 2002. doi: 10.1136/jnnp.73.4.400.

[48] Mari, S., Serrao, M., Casali, C., Conte, C., Martino, G., Ranavolo, A., Coppola, G., Draicchio, F., Padua, L., Sandrini, G., and Pierelli, F. Lower limb antagonist muscle co-activation and its relationship with gait parameters in cerebellar ataxia. The Cerebellum, 13(2):226–236, October 2013. doi: 10.1007/s12311-013-0533-4.

[49] Penn, R.D., Gottlieb, G.L., and Agarwal, G.C. Cerebellar stimulation in man. Journal of Neuro-surgery, 48(5):779–786, May 1978. doi: 10.3171/jns.1978.48.5.0779.

[50] Wolpaw, J.R. What can the spinal cord teach us about learning and memory? The Neuroscientist, 16(5):532–549, October 2010. doi: 10.1177/1073858410368314.

[51] Brownstone, R.M., Bui, T.V., and Stifani, N. Spinal circuits for motor learning. Current Opinion in Neurobiology, 33:166–173, August 2015. doi: 10.1016/j.conb.2015.04.007.

[52] Dimitriou, M. and Edin, B.B. Human muscle spindles act as forward sensory models. Current Biology, 20(19):1763–1767, October 2010. doi: 10.1016/j.cub.2010.08.049.

[53] Wolpert, D.M., Miall, R., and Kawato, M. Internal models in the cerebellum. Trends in Cognitive Sciences, 2(9):338–347, September 1998. doi: 10.1016/s1364-6613(98)01221-2.

[54] Gerwig, M. Timing of conditioned eyeblink responses is impaired in cerebellar patients. Journal of Neuroscience, 25(15):3919–3931, April 2005. doi: 10.1523/jneurosci.0266-05.2005.

[55] Wolf, S.L. and Segal, R.L. Reducing human biceps brachii spinal stretch reflex magnitude. Journal of Neurophysiology, 75(4):1637–1646, April 1996. doi: 10.1152/jn.1996.75.4.1637.

[56] Matthews, P.B. The simple frequency response of human stretch reflexes in which either short-or long-latency components predominate. The Journal of Physiology, 481(3):777–798, December 1994. doi: 10.1113/jphysiol.1994.sp020481.

[57] More, H.L. and Donelan, J.M. Scaling of sensorimotor delays in terrestrial mammals. Proceedings of the Royal Society B: Biological Sciences, 285(1885):20180613, August 2018. doi: 10.1098/r-spb.2018.0613.

[58] Ito, M. Cerebellar microcomplexes. International review of neurobiology, 41:475–487, 1997. doi: 10.1016/s0074-7742(08)60366-9.

[59] d’Avella, A., Fernandez, L., Portone, A., and Lacquaniti, F. Modulation of phasic and tonic mus-cle synergies with reaching direction and speed. Journal of Neurophysiology, 100(3):1433–1454, September 2008. doi: 10.1152/jn.01377.2007.

[60] Marr, D. A theory of cerebellar cortex. The Journal of Physiology, 202(2):437–470, June 1969. doi: 10.1113/jphysiol.1969.sp008820.

[61] Albus, J.S. A theory of cerebellar function. Mathematical Biosciences, 10(1-2):25–61, February 1971. doi: 10.1016/0025-5564(71)90051-4.

[62] Ito, M. Neurophysiological aspects of the cerebellar motor control system. Int. J. Neurol., 7: 126–179, 1970.

[63] Naveros, F., Garrido, J.A., Carrillo, R.R., Ros, E., and Luque, N.R. Event- and time-driven techniques using parallel CPU-GPU co-processing for spiking neural networks. Frontiers in Neuroinformatics, 11, February 2017. doi: 10.3389/fninf.2017.00007.

[64] Mendell, L.M. The size principle: a rule describing the recruitment of motoneurons. Journal of Neurophysiology, 93(6):3024–3026, June 2005. doi: 10.1152/classicessays.00025.2005.

[65] Prochazka, A. Chapter 11 quantifying proprioception. Progress in brain research, 123:133–142, 1999. doi: 10.1016/s0079-6123(08)62850-2.

[66] Al-Falahe, N.A., Nagaoka, M., and Vallbo, A.B. Response profiles of human muscle: afferents during active finger movements. Brain, 113(2):325–346, 1990. doi: 10.1093/brain/113.2.325.

[67] Malik, P., Jabakhanji, N., and Jones, K.E. An assessment of six muscle spindle models for predicting sensory information during human wrist movements. Frontiers in Computational Neuroscience, 9, January 2016. doi: 10.3389/fncom.2015.00154.

[68] Saul, K.R., Hu, X., Goehler, C.M., Vidt, M.E., Daly, M., Velisar, A., and Murray, W.M. Bench-marking of dynamic simulation predictions in two software platforms using an upper limb musculoskeletal model. Computer Methods in Biomechanics and Biomedical Engineering, 18(13): 1445–1458, July 2014. doi: 10.1080/10255842.2014.916698.

[69] Millard, M., Uchida, T., Seth, A., and Delp, S.L. Flexing computational muscle: Modeling and simulation of musculotendon dynamics. Journal of Biomechanical Engineering, 135(2), February 2013. doi: 10.1115/1.4023390.

[70] Delp, S., Loan, J., Hoy, M., Zajac, F., Topp, E., and Rosen, J. An interactive graphics-based model of the lower extremity to study orthopaedic surgical procedures. IEEE Transactions on Biomedical Engineering, 37(8):757–767, 1990. doi: 10.1109/10.102791.

[71] Kreiss, S., Bertoni, L., and Alahi, A. OpenPifPaf: Composite fields for semantic keypoint detection and spatio-temporal association. IEEE Transactions on Intelligent Transportation Systems, 23(8):13498–13511, August 2022. doi: 10.1109/tits.2021.3124981.

[72] Stecina, K., Fedirchuk, B., and Hultborn, H. Information to cerebellum on spinal motor networks mediated by the dorsal spinocerebellar tract. The Journal of Physiology, 591(22):5433–5443, May 2013. doi: 10.1113/jphysiol.2012.249110.

[73] van der Walt, S., Schönberger, J.L., Nunez-Iglesias, J., Boulogne, F., Warner, J.D., Yager, N., Gouillart, E., and Yu, T. scikit-image: image processing in python. PeerJ, 2:e453, June 2014. doi: 10.7717/peerj.453.

[74] Rudolph, K., Axe, M., and Snyder-Mackler, L. Dynamic stability after ACL injury: who can hop? Knee Surgery, Sports Traumatology, Arthroscopy, 8(5):262–269, July 2000. doi: 10.1007/s001670000130.

[75] Li, G., Shourijeh, M.S., Ao, D., Patten, C., and Fregly, B.J. How well do commonly used co-contraction indices approximate lower limb joint stiffness trends during gait for individuals post-stroke? Frontiers in Bioengineering and Biotechnology, 8, January 2021. doi: 10.3389/f-bioe.2020.588908.

[76] Gerstner, W. and Kistler, W.M. Spiking neuron models: Single neurons, populations, plasticity. Cambridge university press, 2002.

[77] Ros, E., Carrillo, R., Ortigosa, E.M., Barbour, B., and Agís, R. Event-driven simulation scheme for spiking neural networks using lookup tables to characterize neuronal dynamics. Neural com-putation, 18(12):2959–2993, 2006. doi: 10.1162/neco.2006.18.12.2959.

